# Class III peroxidases PRX01, PRX44, and PRX73 potentially target extensins during root hair growth in *Arabidopsis thaliana*

**DOI:** 10.1101/2020.02.04.932376

**Authors:** Eliana Marzol, Cecilia Borassi, Philippe Ranocha, Ariel. A. Aptekman, Mauro Bringas, Janice Pennington, Julio Paez-Valencia, Javier Martínez Pacheco, Diana Rosa Rodríguez Garcia, Yossmayer del Carmen Rondón Guerrero, Mariana Carignani, Silvina Mangano, Margaret Fleming, John W. Mishler-Elmore, Francisca Blanco-Herrera, Patricia Bedinger, Christophe Dunand, Luciana Capece, Alejandro D. Nadra, Michael Held, Marisa S. Otegui, José M. Estevez

**Affiliations:** Fundación Instituto Leloir and IIBBA-CONICET. Av. Patricias Argentinas 435, Buenos Aires C1405BWE, Argentina; Université de Toulouse, UPS, UMR 5546, Laboratoire de Recherche en Sciences Végétales, F-31326 CNRS, UMR 5546 Castanet-Tolosan, France; Departamento de Fisiología, Biología Molecular y Celular, Instituto de Biociencias, Biotecnología y Biología Traslacional (iB3). Facultad de Ciencias Exactas y Naturales, Universidad de Buenos Aires, Ciudad Universitaria, Buenos Aires C1428EGA, Argentina; Departamento de Química Biológica, Facultad de Ciencias Exactas y Naturales, Universidad de Buenos Aires (IQUIBICEN-CONICET), Ciudad Universitaria, Buenos Aires C1428EGA, Argentina; Departamento de Química Inorgánica, Analítica y Química Física, Facultad de Ciencias Exactas y Naturales, Universidad de Buenos Aires (INQUIMAE-CONICET), Buenos Aires, CP. C1428EGA, Argentina; Laboratory of Cell and Molecular Biology, University of Wisconsin, Madison, WI, USA; Department of Biology, Colorado State University, Fort Collins, Colorado 80523-1878, USA; Department of Chemistry and Biochemistry, Ohio University, Athens, OH 45701, USA; Centro de Biotecnología Vegetal, Facultad de Ciencias de la Vida, Universidad Andrés Bello and Millennium Institute for Integrative Biology (iBio), Santiago, Chile; Center of Applied Ecology and Sustainability (CAPES), Chile; Departments of Botany and Genetics, University of Wisconsin, Madison, WI, USA

**Keywords:** Arabidopsis, cell walls, extensins, root hairs, ROS, class-III peroxidases

## Abstract

- Root hair cells are important sensors of soil conditions. Expanding several hundred times their original size, root hairs grow towards and absorb water-soluble nutrients. This rapid growth is oscillatory and is mediated by continuous remodelling of the cell wall. Root hair cell walls contain polysaccharides and hydroxyproline-rich glycoproteins including extensins (EXTs).
- Class-III peroxidases (PRXs) are secreted into the apoplastic space and are thought to trigger either cell wall loosening, mediated by oxygen radical species, or polymerization of cell wall components, including the Tyr-mediated assembly of EXT networks (EXT-PRXs). The precise role of these EXT-PRXs is unknown.
- Using genetic, biochemical, and modeling approaches, we identified and characterized three root hair-specific putative EXT-PRXs, PRX01, PRX44, and PRX73. The triple mutant *prx01,44,73* and the PRX44 and PRX73 overexpressors had opposite phenotypes with respect to root hair growth, peroxidase activity and ROS production with a clear impact on cell wall thickness.
- Modeling and docking calculations suggested that these three putative EXT-PRXs may interact with non-*O*-glycosylated sections of EXT peptides that reduce the Tyr-to-Tyr intra-chain distances in EXT aggregates and thereby may enhance Tyr crosslinking. These results suggest that these three putative EXT-PRXs control cell wall properties during the polar expansion of root hair cells.

## Introduction

Primary cell walls, composed by a diverse network containing mainly polysaccharides and a small amount of structural glycoproteins, regulate cell elongation, which is crucial for several plant growth and developmental processes. Extensins (EXTs) belong to hydroxyproline (Hyp)-rich glycoprotein (HRGP) superfamily and broadly include related glycoproteins such as proline-rich proteins (PRPs) and leucine-rich repeat extensins (LRXs) with multiple Ser-(Pro)_3–5_ repeats that may be *O*-glycosylated and contain Tyr (Y)-based motifs (Lamport et al. 2011; Marzol et al. 2018). EXTs require several modifications before they become functional (Lamport et al., 2011; Marzol et al. 2018). After being hydroxylated and *O*-glycosylated in the secretory pathway, the secreted *O*-glycosylated EXTs are crosslinked and insolubilized in the plant cell wall by the oxidative activity of secreted class-III peroxidases (PRXs) on the Tyr-based motifs (Baumberger 2001, 2003; Ringli 2010; Held et al. 2004; Lamport et al., 2011; Chen et al. 2015; Marzol et al. 2018). PRXs are thought to facilitate both intra and inter-molecular covalent Tyr–Tyr crosslinks in EXT networks, possibly through the assembly of triple helices (Velasquez et al. 2015a; Marzol et al. 2018) by generating *iso*dityrosine units (IDT) and pulcherosine, or di-*iso*dityrosine (Di-IDT), respectively (Brady et al., 1996; 1998; Held et al. 2004). In addition, *O*-glycosylation levels in EXTs also affect their insolubilization process in the cell wall (Chen et al. 2015; Velasquez et al. 2015a) since it might influence the EXT interactions with other cell wall components (Nuñez et al., 2009; Valentin et al., 2010). However, the underlying molecular mechanisms of EXT crosslinking and assembly have not been fully determined. It is proposed that *O*-glycosylation levels as well as the presence of Tyr-mediated crosslinking in EXT and related glycoproteins allow them to form a dendritic glycoprotein network in the cell wall. This EXT network affects *de novo* cell wall formation during embryo development (Hall and Cannon 2002; Cannon *et al.*, 2008), they are also implicated in roots, petioles and rosette leaves growth (Saito et al 2014; Møller et al. 2017) and in polar cell expansion processes in root hairs (Baumberger 2001, 2003; Ringli 2010; Velasquez et al. 2011; 2012; 2015a,b) as well as in pollen tubes (Fabrice et al. 2018; Sede et al. 2018; Wang et al. 2018).

Apoplastic class-III PRXs are heme-iron-dependent proteins, members of a large multigenic family in land plants, with 73 members in *Arabidopsis thaliana* (Passardi et al. 2004; Weng and Chapple, 2010). These PRXs catalyze several different classes of reactions. PRX activities coupled to _apo_ROS molecules (_apo_H_2_O_2_) directly affect the degree of cell wall crosslinking (Dunand et al. 2007) by oxidizing cell wall compounds and leading to stiffening of the cell wall through a peroxidative cycle (PC) (Passardi et al. 2004, Cosio & Dunand 2009; Lamport et al. 2011). By constrast, _apo_ROS coupled to PRX activity enhances non-enzymatic cell wall-loosening by producing oxygen radical species (e.g., ^●^OH) and promoting growth in the hydroxylic cycle (HC). In this HC cycle, PRXs catalyze the reaction in which hydroxyl radicals (^●^OH) are produced from H_2_O_2_ after O_2_^●-^ dismutation. In this manner, some PRXs (e.g. PRX36) may function in weaken plant cell walls by the generated ^●^OH that cleave cell wall polysaccharides in seed mucilage extrusion in epidermal cells in the *Arabidopsis* seed coat (Kunieda et al., 2013). It is unclear how these opposite effects on cell wall polymers are coordinated during plant growth (Passardi et al. 2004, Cosio & Dunand 2009; Lee et al. 2013; Ropollo et al. 2011; Lee et al 2018; Francoz et al. 2019). Finally, PRXs also contribute to the superoxide radical (O_2_^●-^) pool by oxidizing singlet oxygen in the oxidative cycle (OC), thereby affecting _apo_H_2_O_2_ levels. Thus, several PRXs are involved in the oxidative polymerization of monolignols in the apoplast of the lignifying cells in xylem (e.g. PRX017, Cosio et al 2017; PRX72, Herrero et al. 2013), in the root endodermis (e.g. PRX64; Lee et al. 2013; Ropollo et al. 2011), and in petal detachment (Lee et al 2018). In addition, PRXs are able to polymerize other components of the plant cell wall such as suberin (Bernards et al., 1999), pectins (Francoz et al. 2019), and EXTs (Schnabelrauch *et al.*, 1996; Jackson et al., 2001). Although several candidates of PRXs have been associated specifically with EXT-crosslinking (EXT-PRXs) by *in vitro* studies (Schnabelrauch et al., 1996; Wojtaszek et al., 1997; Jackson et al., 2001; Price et al., 2003; Pereira et al. 2011; Dong et al., 2015) or based on an immunolabelling extensin study linked to a genetic profile (Jacobowitz et al. 2019), the *in vivo* characterization and mode of action of these EXT-PRXs remain largely unknown. In this work, we used a combination of reverse genetics, molecular and cell biology, computational molecular modeling, and biochemistry to identify three apoplastic PRXs, PRX01, PRX44 and PRX73, as key enzymes possibly potentially involved in Tyr-crosslinking of cell wall EXTs in growing root hair cells. In addition, we propose a hypothetical model in which *O*-glycosylation levels on the triple helixes of EXTs might regulate the degree of Tyr-crosslinking affecting the expansion properties of cell walls as suggested before based on the extended helical polyproline-II conformation state of EXTs (Stafstrom & Staehelin 1986; Owen et al., 2010; Ishiwata et al., 2014) together with an experimental Atomic Force Microscopic (AFM) analysis of crosslinked EXT3 monomers (Cannon et al. 2008) linked to modelling approaches (Velasquez et al. 2015a; Marzol et al 2018). Our results open the way for the discovery of similar interactions in EXT assemblies during root hair development and in response to the environmental changes, such fluctuating nutrient availability in the soil.

## Results and Discussion

In this work, we have chosen to analyze root hair cells because they are an excellent model for tracking cell elongation and identifying PRXs involved in EXT assembly. In previous work, the phenotypes of mutants for PRX01, PRX44 and PRX73 suggested that these PRXs are involved in root hair growth and ROS homeostasis, although their mechanisms of action remained to be characterized (Mangano et al. 2017). All three PRXs are under the transcriptional regulation of the root hair specific transcription factor RSL4 (Yi et al. 2010; Mangano et al. 2017). As expected, these three PRXs are also highly co-expressed with other root hair-specific genes encoding cell wall EXTs (e.g., EXT6-7, EXT12-14, and EXT18) and EXT-related glycoproteins (e.g. LRX1 and LRX2), which functions in cell expansion (Ringli 2010; Velasquez et al. 2011; Velasquez et al. 2015b) (**Figure S1**). Based on this evidence, we hypothesized that these three PRXs might be EXT-PRXs and catalyze Tyr-crosslinks to assemble EXTs in root hair cell walls.

To validate that PRX01, PRX44, and PRX73 are expressed specifically in root hairs, we made transcriptional reporters harboring GFP-tagged fusions of the promoter regions of their genes. In agreement with the *in silico* database (Mangano et al. 2017 and **Figure S1**), all three genes were strongly expressed in root hair cells during cell elongation (**Figure 1A**). Single mutants for these three PRXs showed almost normal root hair growth (Mangano et al. 2017), suggesting a high degree of functional redundancy. Double combinations of *prx44 prx73* (Mangano et al. 2017), *prx01 prx44* and *prx01 prx73* (this study, not shown) as well as the triple null mutant, *prx01 prx44 prx73* showed similarly shorter root hair cells (**Figure 1B**) than what was previously reported for each of the individual *prx* mutants (Mangano et al. 2017). We also obtained two independent lines for each overexpressing PRXs fused to GFP and under the control of a strong *35SCaMV* promoter (PRX^OE^). Unlike the *prx01 prx44 prx73* triple mutant, the lines overexpressing PRX44 and PRX73 had significantly longer root hairs than the Wt Col-0 control (**Figure 2A–B**). The root hairs of the PRX01^OE^ lines, however, were similar to those of Wt Col-0 (**Figure 2A–B**). We reasoned that the lack of enhanced root hair expansion in the PRX01^OE^ lines could be due to reduced levels of overexpression compared to the PRX44^OE^ and PRX73^OE^ lines. However, based on the GFP signals in intact roots (**Figure 2C**), we established that PRX01^OE^ and PRX44^OE^ are strongly expressed, whereas PRX73^OE^ showed more moderate expression. Furthermore, the three PRX-GFP-fusion proteins were detected at the expected molecular weights in an immunoblot (**Figure 2D**), indicating that the tagged proteins are stable. The lack of root hair growth enhancement in PRX01^OE^ line might be due to regulatory aspects on the protein activity rather than in the protein level. Together, these results highlight the partially redundant roles of PRX01, PRX44, and PRX73 as positive regulators of polar growth. This is in agreement with the negative effect of SHAM (salicylic hydroxylamino acid), a peroxidase activity inhibitor (Ikeda-Saito, Shelley et al. 1991; Davey and Fenna 1996), on root hair growth (Mangano et al. 2017). Here is important to highlight that a SHAM treatment produce a more drastic effect on root hair growth and on the inhibition on overall peroxidase activity in the roots (Mangano et al. 2017) than the triple mutant *prx01 prx44 prx73*, suggesting the implication of other unidentified PRXs.

**Figure 1.**
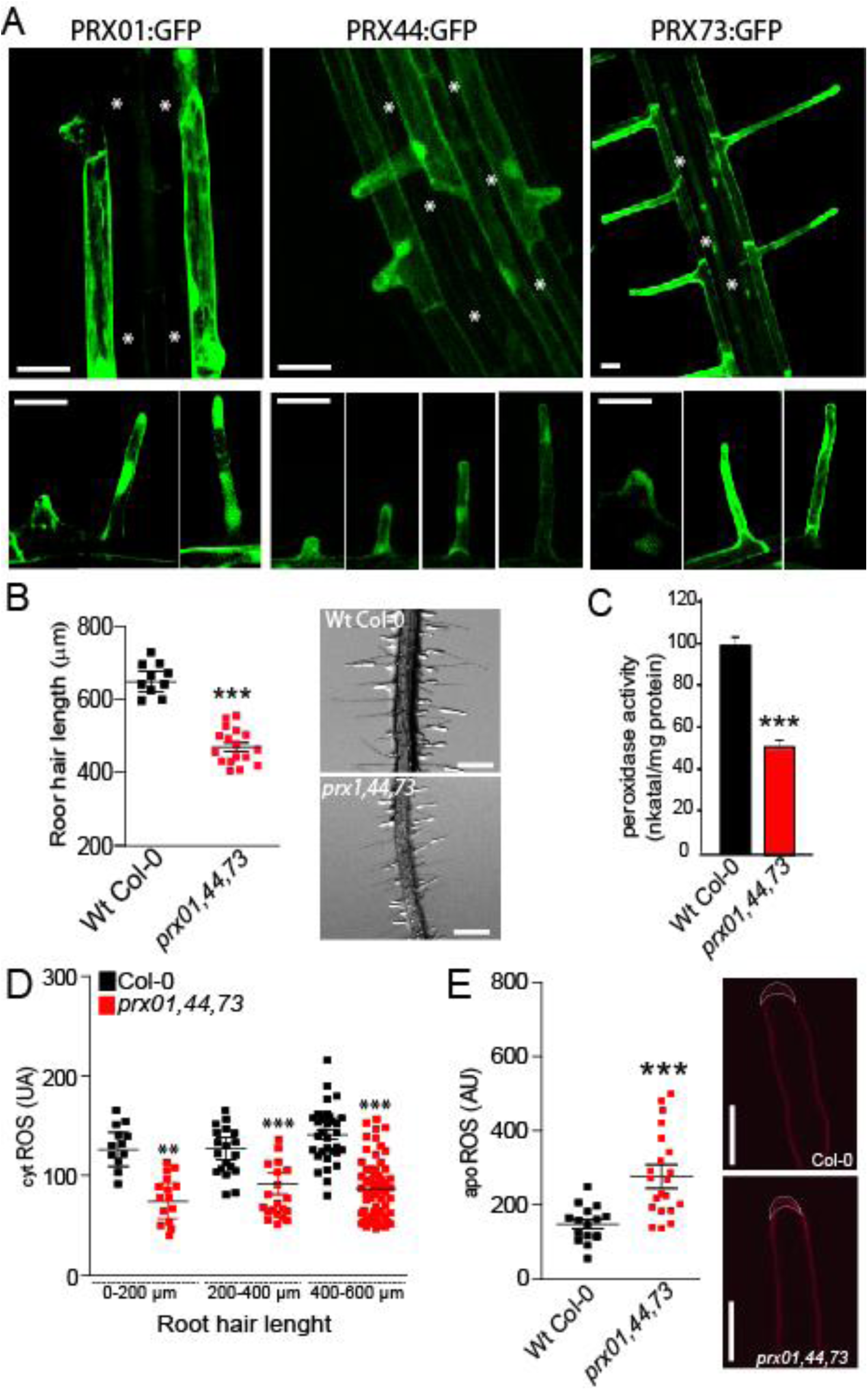
Characterization of root hair-specific PRX01, PRX44 and PRX73 expression and mutant analysis. (**A**) GFP-tagged transcriptional reporters of PRX01, PRX44 and PRX73 show expression in the root elongation zone and specifically in root hairs (bottom). Scale bar = 20 μm. (*) indicates atrichoblast cell layers, which lack GFP expression. (**B**) Root hair length phenotype of Wt and the *prx01,44,73* triple mutant. Left, box-plot of root hair length. Horizontal lines show the means. P-value determined by one-way ANOVA, (***) P<0.001. Right, bright-field images exemplifying the root hair phenotype in each genotype. Scale bars, 1 mm. (**C**) Peroxidase activity in Wt and *prx01,44,73* triple mutant roots. Enzyme activity values (expressed as nkatal/mg protein) are shown as the mean of three replicates ± SD. P-value determined by one-way ANOVA, (***) P<0.001. (**D**) Cytoplasmic ROS levels measured with H_2_DCF-DA in Wt and *prx01,44,73* triple mutant root hairs. Horizontal lines show the means. P-values determined by one-way ANOVA, (***) P<0.001 and (**) P<0.01. (**E**) Apoplastic ROS levels measured with Amplex™ UltraRed (AUR) in Wt and *prx01,44,73 73* triple mutant root hairs. ROS signal was quantified from the root hair cell tip. Left, box-plot of apoROS values. Horizontal lines show the means. P-value determined by one-way ANOVA, (***) P<0.001. Right, fluorescence images exemplifying apoROS detection in root hair apoplast.

**Figure 2.**
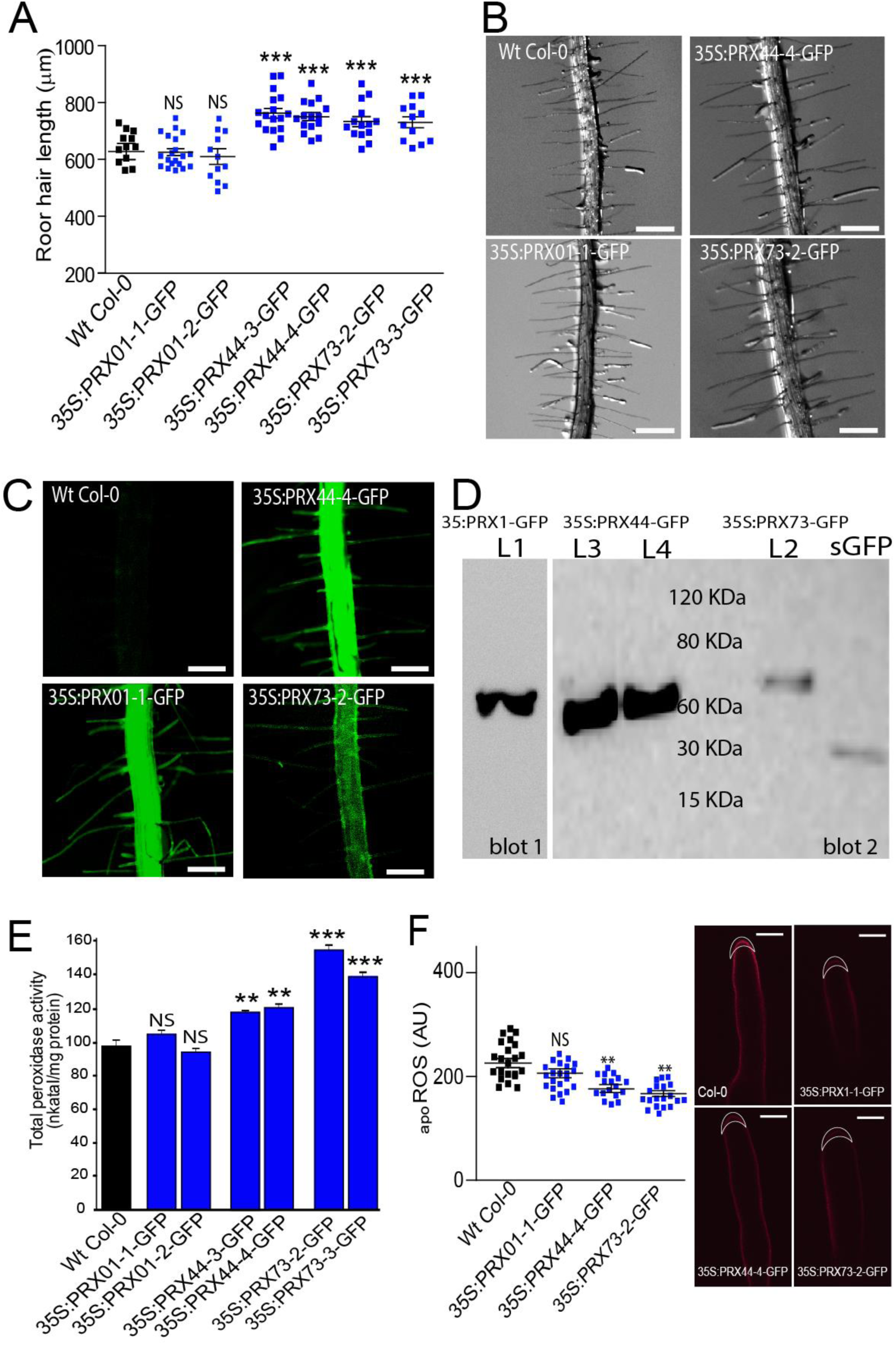
Over-expression of PRX44 and PRX73 promotes root hair growth and higher root peroxidase activity. (**A**) Root hair length phenotype of Wt and PRX^OE^ lines (in Wt background). Box-plot of root hair length. Horizontal lines show the means. P-values determined by one-way ANOVA, (***) P<0.001, (NS) not significantly different. (**B**) Bright-field images exemplifying the root hair phenotype analyzed in Figure 2A. Scale bar = 0.5 mm. (**C**) Expression of GFP-tagged 35S:PRX01, 35S:PRX44 and 35S:PRX73 in root hair cells. (**D**) Western blot of PRX01-GFP, PRX44-GFP and PRX73-GFP. Soluble GFP (sGFP) was used as control. The predicted molecular weights are 62.6 KDa for PRX01-GFP, 60.8 KDa for PRX44-GFP, 62.9 KDa for PRX73 and 27 KDa for sGFP. (**E**) Assays of total peroxidase activity in Wt and PRXs^OE^ lines (in Wt background). Enzyme activity (expressed in nkatal/mg protein) was determined by a guaiacol oxidation-based assay. Values are the mean of three replicates ± SD. P-values determined by one-way ANOVA, (***) P<0.001, (**) P<0.01, (NS) not significantly different. (**F**) Apoplastic ROS levels measured with Amplex™ UltraRed (AUR) in Wt and PRX^OE^ lines (in Wt background). ROS signal was quantified from the root hair cell tip. Left, box plot of apoROS values. Horizontal lines show the means. P-values determined by one-way ANOVA, (**) P<0.01, (NS) not significantly different. Right, fluorescence images exemplifying apoROS detection in root hair apoplast. Scale bar = 10 μm.

To confirm that our mutant and overexpressing lines had the expected changes in peroxidase activity, we measured *in vitro* total peroxidase activity using a guaiacol oxidation-based assay. The *prx01,44,73* roots showed reduced peroxidase activity (close to 50% reduction) (**Figure 1C**), whereas there was a 40–50% increase in PRX73^OE^ and an approximately 20% increase in PRX44^OE^ (**Figure 2E**). Consistent with our root hair growth analysis (**Figure 2A**), PRX01^OE^ showed normal peroxidase activity (**Figure 2E**). The homeostasis and levels of ROS (mostly H_2_O_2_) that regulates polar growth of root hair cells (Mangano et al. 2017) is composed by apoplastic ROS (_apo_ROS) as well as cytoplasmatic ROS pools (_cyt_ROS). Both pools of ROS, their homeostasis and levels are modulated by their transport from the apoplast to the cytoplasmic side by specific aquaporins (PIPs for plasma membrane intrinsic proteins) in plant cells (Dynowski et al., 2008; Hooijmaijers et al., 2012 Rodrigues et al. 2017). We hypothesized that these three PRXs might change the levels of ROS, most probably H_2_O_2_, for their catalytic functions in the cell wall/apoplast. Therefore, we measured _cyt_ROS levels by oxidation of H_2_DCF-DA and _apo_ROS levels with the Amplex Ultra Red (AUR) probe in root hair tips. The *prx01,44,73* root hair tips showed lower levels of _cyt_ROS (**Figure 1D**) but increased _apo_ROS accumulation (**Figure 1E**) compared to Wt Col-0. The _apo_ROS levels were similar in PRX01^OE^, and slightly lower in PRX44^OE^, and PRX73^OE^ lines when compared to Wt Col-0 (**Figure 2F**). These results suggest that PRX01, PRX44, and PRX73 function as apoplastic regulators of ROS-linked root hair cell elongation.

Next, to further analyze the ultrastructure of the cell wall in growing root hairs, we analyzed Wt Col, PRX44^OE^, and *prx01,44,73* triple mutant roots treated or not with SHAM by transmission electron microscopy (**Figure 3A**). Much found thinner cell walls at the root hair tips of PRX44^OE^ (0.74 ± SD 0.24 μm for PRX44^OE^) and *prx01,44,73* (0.61 ± SD 0.14 μm) when compared to Wt Col-0 plants (1.2 ± SD 0.3 μm for Wt) (**Figure 3B**). SHAM treatment caused a statistically significant increase in cell wall thickness in the PRX44^OE^ and *prx01,44,73* root hairs (**Figure 3B**), but not in Wt Col-0. This result suggests the importance of peroxidase activity in cell wall structure and highlights that either depletion of PRX01,44,73 (triple mutant) or the overexpression of PRX44 results in an overall reduction in cell wall thickness in growing root hairs. This implies that the constitutive mis-regulation of PRX activity (either reduced/impaired function or overexpression) affects the capacity of root hairs to form normal cell walls and this clearly affects their cell expansion process.

**Figure 3.**
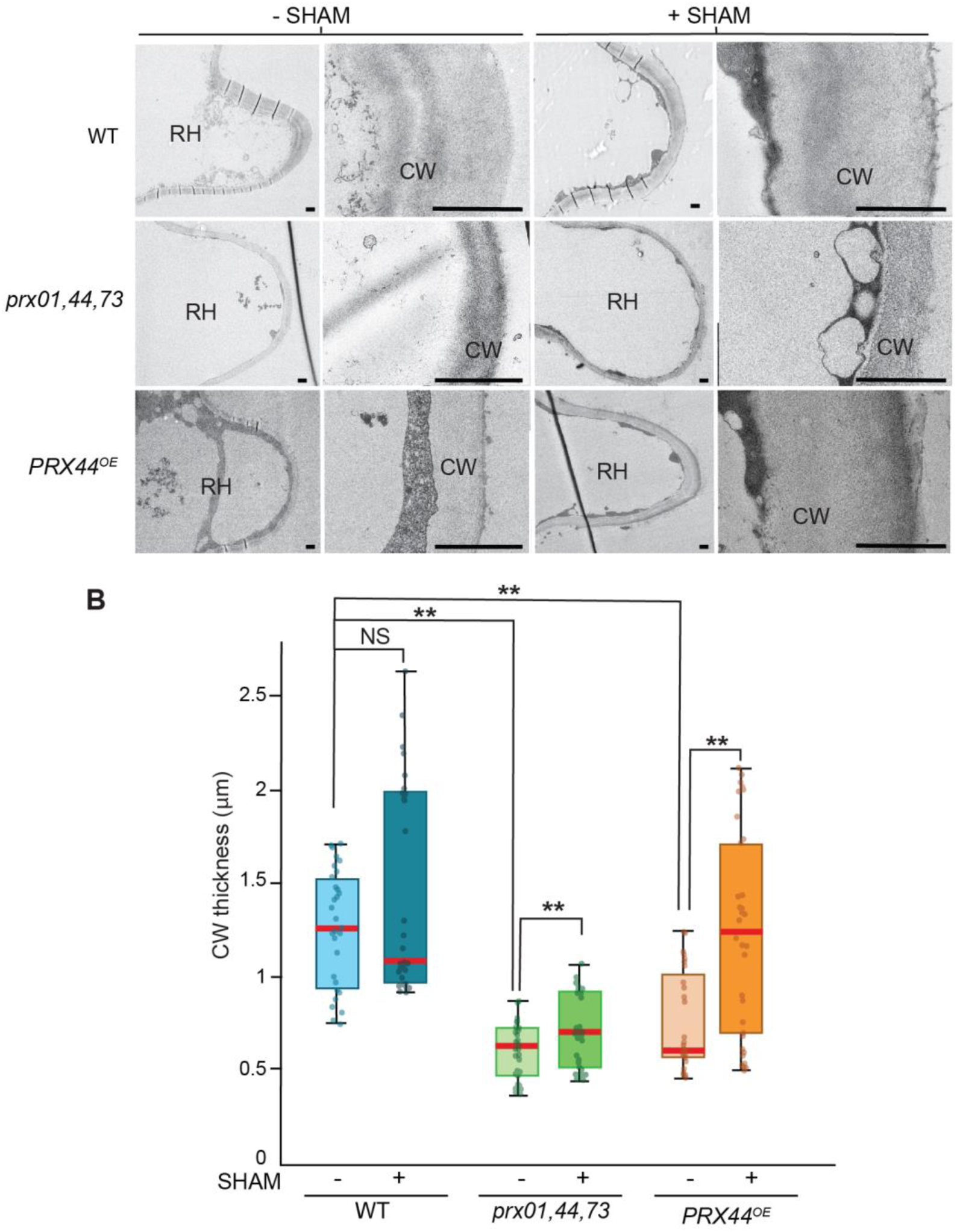
Effect of PRX expression on cell wall thickness in root hair tips. (**A**) Transmission electron micrographs of root hair tips from Wt, *prx1,44,73* triple mutant, and PRX44^OE^ with (+) and without (-) peroxidase inhibitor SHAM. For each genotype and treatment, a representative overview of a root hair (RH) and a detail of the cell wall at the root hair tip (CW) is shown. Scale bar = 1 μm. (**B**) Box and whisker plot showing cell wall thickness measured at the root hair tip of the three genotypes with or without SHAM treatment. (**) P<0.001 determined by t-test. (NS) not significantly different.

Then, we designed an EXT reporter to track EXT secretion and PRX-mediated insolubilization in the cell walls during root hair cell elongation. The secreted EXT reporter carries a Tomato tag (SS-TOM-Long-EXT) that is fluorescent under the acidic pH (Shen et al. 2014) that is typical of plant cell walls and apoplastic spaces (Stoddard & Rolland 2018). A secreted Tomato tag (ss-TOM) was used as a control (**Figure S2A**). The EXT domain includes only two Tyr, which are at the C-terminus and separated by 10 amino acids (Stratford et al., 2001). Expression of the EXT reporter was first tested in onion (*Allium cepa*) cells, and then the reporter was stably expressed in Arabidopsis root hairs (**Figure S2C-F**). In both cases, plasmolysis was used to retract the plasma membrane from the cell surface to show that the EXT reporter was localized in the cell walls. Using immunoblot analysis, we detected the full-length EXT-Tomato fusion protein, with possible *O*-glycan modifications, running as higher molecular weight bands than expected (**Figure S2B**). Importantly, the EXT reporter did not interfere with the polar growth of root hairs (**Figure S2D**), and, therefore, could be used to track changes in the *in situ* arrangement of cell wall EXTs. SS-TOM-Long-EXT is clearly secreted in the cell wall of growing root hairs (**Figure S2C**) but remains to be tested if these EXT reporter is mislocalized under an inhibited PRX environment (SHAM treated) or in *prx01,44,73* mutant background.

We then assessed the level of crosslinking of EXT Tyr residues by measuring peptidyl-tyrosine (Tyr) and isodityrosine (IDT, dimerized Tyr) in EXT extracted from whole roots. We detected a significant increase in peptidyl-Tyr in the *prx01,44,73* triple mutant relative to Wt Col-0, and slightly higher levels of IDT in EXTs extracted from the PRX73^OE^ line (**Table 1**). By contrast, we identified strong downregulation of Tyr- and IDT-levels in the EXT under-*O*-glycosylation mutants *p4h5 sergt1-1*, and *sergt1-1 rra3* (**Table 1**). In these two double mutants, root hair growth is drastically inhibited (Velasquez et al. 2015a). PROLYL 4-HYDROXYLASE (P4H5), PEPTIDYL-SER GALACTOSYLTRANSFERASE (SGT1/SERGT1), and REDUCE RESIDUAL ARABINOSE 3 (RRA3) are key enzymes that modify EXT hydroxylation (P4H5) and EXT *O*-glycosylation (SERGT1 and RRA3) (Marzol et al. 2018). Specifically, it was shown that P4H5 is a 2-oxoglutarate (2OG) dioxygenase that catalyze the formation of trans-4-hydroxyproline (Hyp/O) from peptidyl-proline preferentially in an EXT context allowing these proteins to be *O*-glycosylated (Velasquez et al. 2011; Velasquez et al. 2015b). In the case of RRA3, together with RRA1–RRA2 homologous proteins (Egelund et al., 2007; Velasquez et al., 2011), they are thought to transfer the second arabinose to the sort glycan (composed by 4–5 units of L-arabinofuranose) attached to the Hyp in the EXT peptides. SERGT1 add the single galactose units to the serine in the repetitive motif of Ser-(Pro)_3–5_ present in EXT and EXT-related proteins (Saito et al. 2014). These results are consistent with the notion that *O*-glycans strongly affect EXT Tyr crosslinking, as was previously suggested based on the drastically reduced root hair growth of the under-glycosylation mutants and *in vitro* crosslinking rates (Velasquez et al 2015a,b; Chen et al. 2015). We hypothesize that absent or low *O*-glycosylation of EXTs or an increase in PRX levels may trigger a reduction in the amount of peptidyl-Tyr and IDT levels in EXTs, with a putative concomitant increase in the amounts of higher-order Tyr crosslinks (trimers as Pulcherosine and tetramers as Di-IDT), thus inhibiting root hair growth. For technical reasons we could not measure the Pulcherosine and Di-IDT levels described before in EXTs (Brady et al., 1996; 1998; Held et al. 2004) to test this hypothesis. Further research is needed to decipher the *in vivo* regulation of Tyr crosslinking of EXTs by these three PRXs in plant cells.

**Table 1.** Peptidyl-Tyr and iso-dityrosine (IDT) contents in cell walls isolated from Wt, *prx01,44,73* triple mutant, PRX^OE^ lines and mutant lines with under-glycosylated EXTs. P-values were determined by one-way ANOVA, (***) P<0.001, (**) P<0.01. STD=Standard Deviation. Values significantly different than Wt are highlighted in blue if higher and in light blue if lower than Wt Col-0.

|  | ng Tyr/μg CW (STD) | ng IDT/μg CW (STD) |
| --- | --- | --- |
| Wt Col-0 | 7.799 ± 0.26 | 0.853 ± 0.08 |
| *prx01 prx44 prx73* | 9.588 ± 0.31** | 0.963 ± 0.02 |
| PRX44<sup>OE</sup> | 8.649 ± 0.07 | 0.953 ± 0.04 |
| PRX73<sup>OE</sup> | 8.700 ± 0.12 | 1.042 ± 0.02** |
| *under O-glycosylated EXTs* |  |  |
| *sergt1-1 rra3* | 3.530 ± 0.08*** | 0.235 ± 0.01*** |
| *p4h5 sergt1-1* | 3.766 ± 0.06*** | 0.225 ± 0.02*** |

A major limitation in our understanding of how EXTs function in plant cell walls is the lack of a realistic full-length EXT protein model. We used coarse-grained molecular dynamics to build a larger model of a triple-helix EXT sequence, that includes 10 conserved repeats (SPPPPYVYSSPPPPYYSPSPKVYYK, 250 aminoacids in each polypeptide chain) (**Figure S3A–B**). Parameters for the *O*-glycosylated form of EXT were developed in this work (**Figure S4**). The EXT molecules were modeled in two different states: as a non-glycosylated trimeric helical conformation similar to animal collagen and in the O-glycosylated state, with 4 arabinose monosaccharides in each hydroxyproline. Those two states were simulated restraining both ends of the polypeptide chains, to model a fully extended helix (consistent with an “indefinitely long-EXT”), and without that restriction, to evaluate the conformation that an isolated 10-repeat triple helix would adopt. The results indicate the importance of the triple-helix conformation in the overall stability of the protein and especially in the conservation of its fibril-like structure, in agreement with shorter-repeats single helix simulations performed previously (Velasquez et al. 2015a; Marzol et al. 2018). The total volume of the extended systems triple helix was measured in both glycosylation states (**Table S1**), differentiating EXT-protein-only and EXT-protein+glycan volumes for the fully *O*-glycosylated EXT state. We observed that the EXT-protein-only volume was significantly augmented by the presence of the oligosaccharide moieties, indicating that O-glycans increase the distance between peptide chains in the EXT triple helix. We report the average diameters for those systems (**Table S1**), which are consistent with the diameters previously reported based on Atomic Force Microscopy (AFM) (images (Cannon et al. 2008). Additionally, *O*-glycosylation contributes to an increase in the average distance between the side chains of tyrosine residues, decreasing the proportion of tyrosine side chains that are close enough to lead to crosslinked EXT chains (**Figure S3C**). Current experimental and modeling lines of evidence are in agreement with the proposed role of proline-hydroxylation and carbohydrate moieties in keeping the EXT molecule in an extended helical polyproline-II conformation state (Stafstrom & Staehelin 1986; Owen et al., 2010; Ishiwata et al., 2014). This extended conformation might allow EXTs to interact properly with each other and with other components in the apoplast, including PRXs and pectins, to form a proper cell wall network (Nuñez et al., 2009; Valentin et al., 2010).

To test if these three PRXs (PRX01, PRX44, and PRX73) might be able to interact with single-chain EXTs, we performed homology modeling with GvEP1, an EXT-PRX that is able to crosslink EXTs *in vitro* (Jackson et al., 2001; Pereira et al. 2011). In addition, we included PRX64, as a PRX described for lignin polymerization in the root endodermis (Lee et al. 2013) and PRX36, which is able to bind homogalacturonan pectin in the seed coat (Francoz et al. 2019) as controls. By *docking* analysis, we obtained interaction energies (Kcal/mol) for all of them. We analyzed docking with four different short EXT peptides: a non-hydroxylated peptide, a hydroxylated peptide, an arabinosylated peptide and an arabino-galactosylated peptide. As mentioned earlier, it was previously shown that mutants carrying under-*O*-glycosylated EXTs have severe defects in root hair growth (Velasquez et al. 2011; Velasquez et al. 2015a). Our docking results for the different PRXs show consistent interaction energy differences that depend on the EXT glycosylation state, being higher for non-*O*-glycosylated species. In addition, *O*-glycosylated EXT variants docked in a rather dispersed way while non-*O*-glycosylated variants preferentially docked in a grooved area (**Figure 4A–C**). Furthermore, **Figure 4A** shows how a non-*O*-glycosylated peptide binds through a groove, leaving one Tyr docked in a cavity and very close to the heme iron (5Å), with a second Tyr a few Angstroms away. The arrangement and distances between the tyrosines suggest that this could be an active site where Tyr crosslinking takes place. Although it is not possible to compare the interaction energies obtained with the different EXT species among docking runs, a general trend can be observed in **Figure 4C**. In general, we observed higher interaction energies (more negative values) for hydroxylated EXT species, followed by non-hydroxylated EXTs, and then by *O*-glycosylated EXT variants. When we compared interaction energies among different PRXs interacting with EXT substrates with the same degree of *O*-glycosylation, we observed that PRX73 displayed the highest interaction activity with the non-hydroxylated EXT species, followed by PRX01 and then PRX44. For the hydroxylated EXT variant, the order was PRX44>PRX73>PRX01. PRX44 displayed the highest interaction energy with the *O*-glycosylated species. All together, these results are consistent with the constitutive root hair growth effect observed for PRX44^OE^ and PRX73^OE^ and with non-glycosylated EXT being the substrate of peroxidation. Overall, this possibly indicates that PRX44 and PRX73 might interact with EXT substrates and possibly catalyze Tyr-crosslinking in open regions of the EXT backbones with little or no *O*-glycosylation. This is in agreement with previous studies that suggested that high levels of *O*-glycosylation in certain EXT segments physically restrict EXT lateral alignments, possibly by acting as a branching point (Cannon et al.2008; Velasquez et al., 2015a; Marzol et al. 2018).

**Figure 4.**
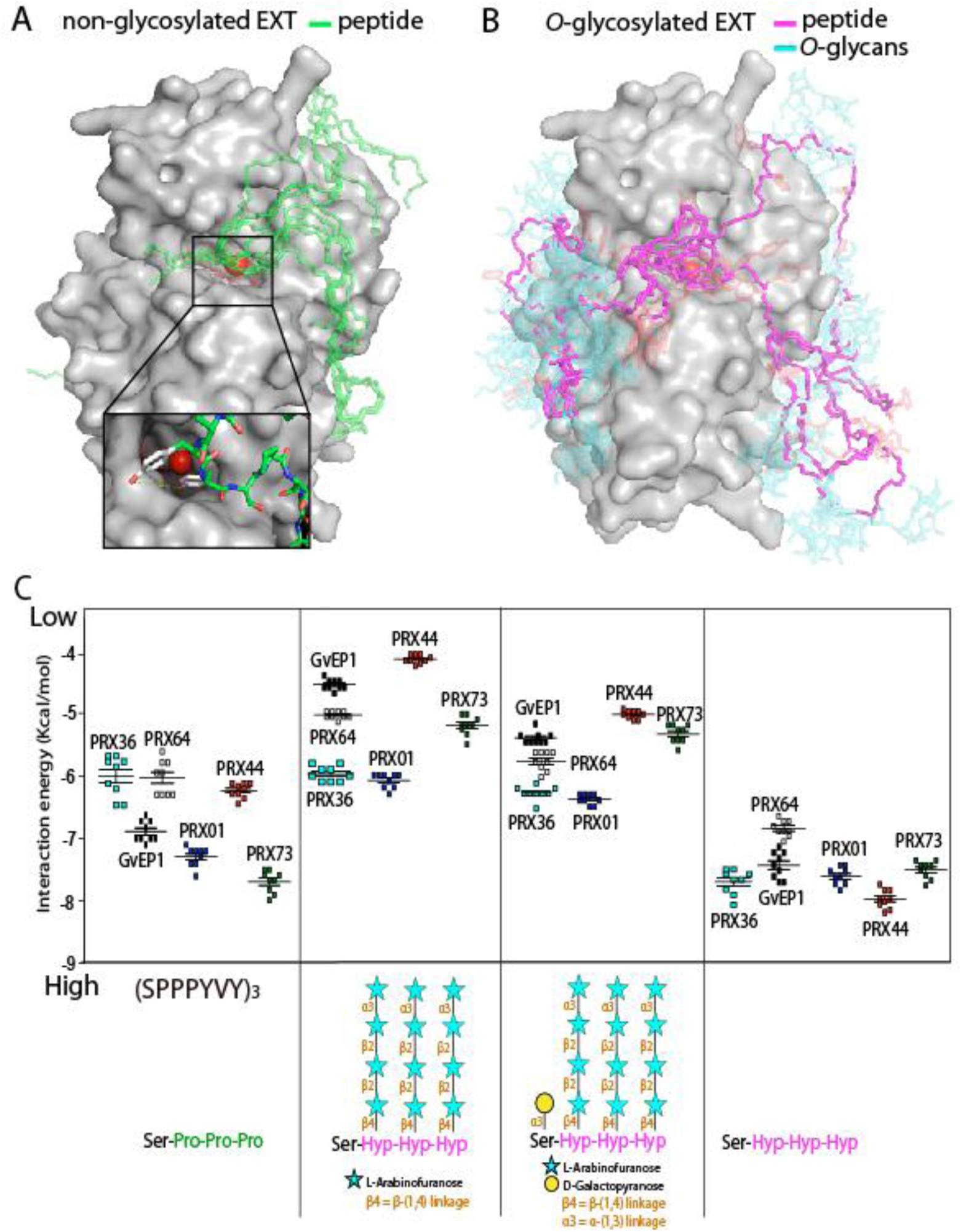
Interaction by an *in silico* docking approach of PRX01, PRX44 and PRX73 with EXT peptides. (**A**,**B**) Ten docking results for each EXT *O*-glycosylation state are shown superimposed on the PRX44 protein surface to evaluate the consistence of docking sites. (**A**) Model of PRX44 (protein surface shown in gray) complexed to a non-*O*-glycosylated EXT substrate (SPPPYVY)_3_ (in green, depicted as sticks). Heme is depicted as thin sticks while iron is a red sphere. Bottom inset, two close tyrosine residues dock near to the possible active site of PRX44. (**B**) Model of PRX44 (protein surface shown in gray) complexed to an *O*-glycosylated-EXT substrate (protein backbone shown in magenta, and *O*-glycans shown in light blue, both depicted as sticks). Heme is depicted as thin sticks while iron is a red sphere. Arabino-galactosylated EXT peptide = [(SOOOYVY)_3_-AG]. (**C**) Comparison of the binding energy of different peroxidases to EXT substrates with different degrees of *O*-glycosylation. A non-hydroxylated EXT peptide (SPPPYVY)_3_, a hydroxylated but not *O*-glycosylated EXT peptide [(SOOOYVY)_3_; O=hydroxyproline], only arabinosylated EXT-peptide [(SOOOYVY)_3_-A], and arabino-galactosylated EXT peptide [(SOOOYVY)_3_-AG] were analyzed.

To examine the evolution of PRX01, PRX44, and PRX73, we performed comprehensive phylogenetic analyses of Class-III peroxidases across diverse land plant lineages. Under low selective pressure to maintain substrate specificity, EXT-PRX activities might have evolved multiple times in parallel during land plant evolution through gene duplication followed by neofunctionalization or subfunctionalization. PRX01, PRX44, and PRX73 belong to three independent orthologous groups (**Figure S5**) and orthologs for each *A. thaliana* PRX have been detected in available Brassicaceae genomes and in various Angiosperm and Gymnosperm families, but not from Lycophytes and from non-vascular land plants. Thus, these three PRX sequences were the result of ancestral duplications before the divergence between Gymnosperms and Angiosperms but after the emergence of the Tracheophytes (**Figure S5**). Orthologs of the three PRX genes have only been detected in true root containing organisms and these three PRXs are expressed in roots and root hairs, as are most of their orthologous sequences (where expression data are available) (**Figure S6**). This strongly supports the hypothesis that the three independent orthogroups have conserved functions in roots. With the exception of PRX73, which belongs to a cluster containing the putative EXT-PRX from tomato (*Solanum lycopersicum*; LePRX38), the other two PRX sequences did not cluster with sequences already described as putative EXT-PRXs, such as PRX09 and PRX40 (Jacobowitz et al. 2019). Indeed, the other known EXT-PRXs (identified mostly based on *in vitro* evidence) are not clustered together, but are widely distributed in the tree (**Figure S5**). This analysis suggests that plant EXT-PRXs might have evolved several times in parallel during Tracheophyte evolution.

Based on the results shown in this work, we propose a working model in which PRX01, PRX44, and PRX73 (and possibly other PRXs) control root hair growth by channelling H_2_O_2_ consumption and affecting the cell wall hardening process. In this polar growing cells, it is known that H_2_O_2_ is primary derived from by the respiratory burst oxidase homolog C (RBOHC), and to a lower extent from RBOHH and RBOHJ activities that produce superoxide ions (Monshauser et al. 2007; Tajeda et al. 2008; Mangano et al. 2017) that are further converted chemically or enzymatically to H_2_O_2_. Then, part of H_2_O_2_ might be transported from the apoplast to the cytoplasm side by specific PIPs as it was shown to occur in several plant cell types (e.g. in stomata and epidermal cells) in response to diverse stimuli (Dynowski et al., 2008; Hooijmaijers et al., 2012; Rodrigues et al. 2017). When apoplastic PRX protein levels are low, which is linked to reduced peroxidase activity as in the triple mutant *prx01,44,73*, high levels of H_2_O_2_ accumulate in the apoplast, triggering through the oxidative cycle (OC) a cell wall loosening effect that affects growth homeostasis and inhibits expansion by decreasing root hair growth and cell wall thickness (**Figure S7**). Concomitantly, deficient PRX activity in the apoplast also triggers lower H_2_O_2_ levels in the cytoplasm of growing root hairs. This is in agreement with the fact that exogenously supplied H_2_O_2_ inhibited root hair polar expansion, whereas treatment with ROS scavengers (e.g., ascorbic acid) caused root hair bursting (Orman-ligeza et al. 2016), reinforcing the notion that _apo_ROS modulates cell growth by impacting cell-wall properties (Mangano et al. 2017). Our results suggest that either low or high levels of apoplastic Class-III PRXs in the root hair cell walls might affect the homeostasis of ROS and cell wall thickness with a clear effect on cell expansion. Still several aspects of this model proposed here remains to be tested.

## Conclusions

Currently, several of the 73 apoplastic Class-III PRXs in *Arabidopsis thaliana* have no assigned biological function. In this work, we have characterized three related EXT-PRXs, PRX01, PRX44, and PRX73 that function in ROS homeostasis and potentially in EXT assembly during root hair growth. These PRXs might control Tyr crosslinking in EXTs and related glycoproteins and modify its secretion and assembly in the nascent tip cell walls. Using modeling and docking approaches, we were able to measure the interactions of these PRXs with single chain EXT substrates. All these lines of evidence indicate that PRX01, PRX44, and PRX73 are important enzymes that could be involved in EXT assembly during root hair growth. From an evolutionary perspective, all the putative EXT-PRXs (previously identified based on *in vitro* evidence or immunolabeling) do not cluster together in the phylogenetic tree of Class-III PRXs, suggesting that plant-related EXT-PRXs might have evolved several times in parallel during Tracheophyte evolution. Interestingly, as a convergent evolutionary extracellular assembly, hydroxyproline-rich collagen Class-IV, similar to the green EXT linage and related glycoproteins, is also crosslinked by the activity of a specific class of animal heme peroxidases (named peroxidasin or PXDN) to form insoluble extracellular networks (Vanacore et al. 2009; Bhave et al. 2012). While the biophysical properties of collagen IV allow the correct development and function of multicellular tissues in all animal phyla (Brown et al. 2017), EXT assemblies also have key functions in several plant cell expansion and morphogenesis processes (Baumberger 2001, 2003; Hall and Cannon et al. 2002; Cannon *et al.*, 2008; Ringli 2010; Lamport et al., 2011; Velasquez et al. 2015a,b; Fabrice et al. 2018; Sede et al. 2018; Marzol et al. 2018). This might imply that crosslinked extracellular matrices based on hydroxyproline-rich polymers (e.g., collagens and EXTs) have evolved more than once during eukaryotic evolution, providing mechanical support to single and multiple cellular tissues. Further analyses are required to establish how these described EXT-PRXs catalyze Tyr crosslinks on EXTs at the molecular level and how this assembly process is regulated during polar cell expansion.

## Experimental Procedures

### Plant and growth conditions

*Arabidopsis thaliana* Columbia-0 (Col-0) was used as the wild Class (Wt) genotype in all experiments. All mutants and transgenic lines tested are in this genetic background. Seedlings were germinated on agar plates in a Percival incubator at 22°C in a growth room with 16h light/8h dark cycles for 10 days at 140 μmol m^-2^s^-1^ light intensity. Plants were transferred to soil for growth under the same conditions. For identification of T-DNA knockout lines, genomic DNA was extracted from rosette leaves. Confirmation by PCR of a single and multiple T-DNA insertions in the target PRX genes were performed using an insertion-specific LBb1.3 primers in addition to one gene-specific primer. To ensure gene disruptions, PCR was also run using two gene-specific primers, expecting bands corresponding to fragments larger than in WT. We isolated homozygous lines for PRX01 (AT1G05240, *prx01-2*, Salk_103597), PRX44 (AT4G26010, *prx44-2*, Salk_057222) and PRX73 (AT5G67400, *prx73-3*, Salk_009296). SERGT1 (*sergt1-1* SALK_054682), *rra3* (GABI_233B05) (Velasquez et al., 2011) and *p4h5* T-DNA mutant (Velasquez et al., 2011) were isolated and described previously. Double and triple mutants were generated by manual crosses of the corresponding single mutants (Velasquez et al., 2015a). All the mutant lines used in this study are described in **Table S2**.

### PRX::GFP and 35S::PRX-GFP lines

Vectors based on the Gateway cloning technology (Invitrogen) were used for all manipulations. Constitutive expression of PRXs-GFP tagged lines were achieved in plant destination vector pMDC83. cDNA PRXs sequences were PCR-amplified with AttB recombination sites. PCR products were then recombined first in pDONOR207 and transferred into pGWB83. To generate transcriptional reporter, the PRXs promoter regions (2Kb) was amplified and recombined first in pDONOR207 and transferred into pMDC111. All the transgenic lines used in this study are described in **Table S2**.

### SS-TOM and SS-TOM-Long-EXT constructs

The binary vector pART27, encoding tdTomato secreted with the secretory signal sequence from tomato polygalacturonase and expressed by the constitutive CaMV 35S promoter (pART-SS-TOM), was the kind gift of Dr. Jocelyn Rose, Cornell University. The entire reporter protein construct was excised from pART-SS-TOM by digesting with *Not*I. The resulting fragments were gel-purified with the QIAquick Gel Extraction Kit and ligated using T4 DNA Ligase (New England Biolabs) into dephosphorylated pBlueScript KS+ that had also been digested with *Not*I and gel-purified to make pBS-SS-TOM. The plasmid was confirmed by sequencing with primers 35S-FP (5’-CCTTCGCAAGACCCTTCCTC-3’) and OCS-RP (5’-CGTGCACAACAGAATTGAAAGC-3’). The sequence of the EXT domain from *SlPEX1* (NCBI accession AF159296) was synthesized and cloned by GenScript into pUC57 (pUC57-EXT). The plasmid pBS-SS-TOM-Long-EXT was made by digesting pUC57-EXT and pBS-SS-TOM with *Nde*I and *SgrA*I, followed by gel purification of the 2243 bp band from pUC57-EXT and the 5545 bp band from pBS-SS-TOM, and ligation of the two gel-purified fragments. The pBS-SS-TOM-Long-EXT plasmid was confirmed by sequencing with 35S-FP, OCS-RP, and tdt-seq-FP (5’-CCCGTTCAATTGCCTGGT-3’). Both pBS plasmids were also confirmed by digestion. The binary vector pART-SS-TOM-Long-EXT was made by gel purifying the *Not*I insert fragment from the pBS-SS-TOM-Long EXT plasmid and ligating it with pART-SS-TOM backbone that had been digested with *Not*I, gel purified, and dephosphorylated. This plasmid was confirmed by sequencing. The construct SS-TOM and SS-TOM-Long-EXT where transformed into *Arabidopsis* plants. The secretory sequence (SS) from tomato polygalacturonase is <u>MVIQRNSILLLIIIFASSISTCRSGT</u> (2.8kDa) and the EXT-Long domain sequence with six alanine cluster is B<u>AAAAAAA</u>CTLPSLKNFTFSKNIFESMDETCRPSESKQVKIDGNENCLGGRSEQRTEKECFPVVSKPVDCSKGHCG VSREGQSPKDPPKTVTPPKPSTPTTPKPNPSPPPPKTLPPPPKTSPPPPVHSPPPPPVASPPPPVHSPPPPVASPPPPVHSPPPPPVASPPPPVHSPPPPVASPPPPVHSPPPPVHSPPPPVASPPPPVHSPPPPVHSPPPPVHSPPPPVHSPPPPVHSPPPPVASPPPPVHSPPPPVHSPPPPVHSPPPPVASPPPPVHSPPPPPPVASPPPPVHSPPPPVASPPPPVHSPPPPVASPPPPVHSPPPPVHSPPPPVHSPPPPVASPPPALVFSPPPPVHSPPPPAPVMSPPPPTFEDALPPTLGSLYAS PPPPIFQGY*395-(39.9kDa). The predicted molecular size for SS-TOM protein is 54.2 kDa and for SS-TOM-EXT-Long Mw is 97.4 kDa. All the transgenic lines used in this study are described in **Table S2**.

### Root hair phenotype

For quantitative analysis of root hair phenotypes in *prx01,44,73* mutant, 35S:PRX-GFP lines and Wt Col-0, 200 fully elongated root hairs were measured (n roots= 20-30) from seedlings grown on vertical plates for 10 days. Values are reported as the mean ±SD using the Image J software. Measurements were made after 7 days. Images were captured with an Olympus SZX7 Zoom microscope equipped with a Q-Colors digital camera.

### Confocal imaging

Root hairs were ratio imaged with the Zeiss LSM 710 laser scanning confocal microscope (Carl Zeiss) using a 40X oil-immersion, 1.2 numerical aperture. EGFP (473–505nm) emission was collected using a 458-nm primary dichroic mirror and the meta-detector of the microscope. Bright-field images were acquired simultaneously using the transmission detector of the microscope. Fluorescence intensity was measured in 7 µm ROI (Region Of Interest) at the root hair apex.

### Peroxidase activity

Soluble proteins were extracted from roots grown on agar plates in a Percival incubator at 22°C in a growth room for 10 days at 140 μmol m^-2^s^-1^ light intensity by grinding in 20mM HEPES, pH 7.0, containing 1 mM EGTA, 10mM ascorbic acid, and PVP PolyclarAT (100mg g^-1^ fresh material; Sigma, Buchs, Switzerland). The extract was centrifuged twice for 10 min at 10,000 g. Each extract was assayed for protein levels with the Bio-Rad assay (Bio-Rad). PRX activity was measured at 25°C by following the oxidation of 8 mM guaiacol (Fluka) at 470 nm in the presence of 2 mM H_2_O_2_ (Carlo Erba) in a phosphate buffer (200 mM, pH6.0). Values are the mean of three replicates ± SD.

### Cytoplasmic ROS (cytROS) measurements

2′,7′-dichlorodihydrofluorescein diacetate (H_2_DCF-DA) is as a cell-permeable fluorogenic probe to quantify reactive oxygen species (ROS). H_2_DCFDA diffuses into cells and is deacetylated by cellular esterases to form 2′,7′-dichlorodihydrofluorescein (H_2_DCF). In the presence of ROS, predominantly H_2_O_2_, H_2_DCF is rapidly oxidized to 2′,7′-dichlorofluorescein (DCF), which is highly fluorescent, with excitation and emission wavelengths of 498 and 522 nm, respectively. To measure cytoplasmic ROS in root hairs cells, growth of Arabidopsis seeds on a plate was done with 1% sterile agar for 8 d in a chamber at 22°C with continuous light. These seedlings were incubated in darkness on a slide for 10 min with 50 μM H_2_DCFDA at room temperature. Samples were observed with Zeiss Imager A2 Epifluorescence. A 10× objective was used, 0.30 N.A., and exposure time 80-500ms. Images were analyzed using ImageJ 1.50b software. To measure ROS mean, a circular region of interest (ROI) (r=2.5) was chosen in the tip zone of the root hair. All root hairs of six seedlings per genotype were analyzed. The reported values are the mean ± standard deviation (mean ± SD).

### Apoplastic ROS (_apo_ROS) measurements

To measure apoplastic ROS in root hair cells, roots of 7-day-old seedlings were incubated with 50 µM Amplex™ UltraRed Reagent (AUR, Molecular Probes) for 20 min in dark conditions and rinsed with liquid MS. Root hairs were imaged with a Zeiss LSM5 Pascal laser scanning confocal microscope. The fluorescence emission of oxidized AUR in the apoplast of root hair cells was observed between 585 and 610 nm using 543 nm argon laser excitation, 40X objective, N/A=1.2. The intensity of fluorescence was quantified on digital images using ImageJ software. Quantification of the AUR probing fluorescence signal was restricted to apoplastic spaces at the root hair tip (as shown in **Figure 1**). The measurements were performed in three independent experiments (n = 6) with the same microscopic settings.

### Phylogenetic analysis

73 class-III PRX protein sequences from *A. thaliana*, two putative lignin class-III PRXs from *Zinnia elegans* and 4 putative Extensin class-III PRXs from *Lupinus album, Lycopersicum esculentum, Phaseolus vulgaris* and *Vitis vinifera*, have been aligned with ClustalW and the tree constructed using the Neighbor-Joining method (Saitou and Nei, 1987). The analyses were conducted in MEGA7 (Kumar, 2016). All protein sequences are available using their ID number (http://peroxibase.toulouse.inra.fr (Savelli et al., 2019).

### Co-expression analysis network

Co-expression networks for *RSL4* root hair genes were identified from PlaNet (http://aranet.mpimp-golm.mpg.de) and trimmed to facilitate readability (Mutwill et al. 2011). Each co-expression of interest was confirmed independently using the expression angler tool from Botany Array Resource BAR (http://bar.utoronto.ca/ntools/cgi-bin/ntools_expression_angler.cgi) and ATTED-II (http://atted.jp). Only those genes that are connected with genes of interest are included.

### Tyr-crosslinking analysis

Alcohol-insoluble residues of root tissues obtained from *PRX01,44,73* mutants, Col-0 and 35Sp::PRX44-GFP lines were hydrolyzed in 6 N HCl (aqueous) with 10 mM phenol (2 mg ml^-1^; 110 °C; 20 h). Hydrolysates were dried under a steady stream of nitrogen (gas) and then re-dissolved at 10 mg ml^-1^ in water. The hydrolysates were fractionated by gel permeation chromatography on a polyhydroxyethyl A column (inner diameter, 9.4 x 200 mm, 10 nm pore size, Poly LC Inc., Columbia, MD) equilibrated in 50 mM formic acid and eluted isocratically at a flow rate of 0.8 ml min^-1^. UV absorbance was monitored at 280 nm. The amounts of Tyr and IDT in the hydrolysates were then determined by comparison with peak areas of authentic Tyr and IDT standards. Response factors were determined from three level calibrations with the Tyr and IDT standards.

### Immuno-blot Analysis

Plant material (100 mg of root from 15 days old seedlings grown as indicated before) was collected in a microfuge tube and ground in liquid nitrogen with 400 mL of protein extraction buffer (125 mM Tris-Cl, pH. 4.4, 2% [w/v] SDS, 10% [v/v] glycerol, 6M UREA, 1% [v/v] b-mercaptoethanol, 1mM PMSF). Samples were immediately transferred to ice. After 4° centrifugations at 13000 rpm for 20 min, supernatant was move to a new 1.5 ml tube and equal volumes of Laemmli buffer (125 mM Tris-Cl, pH. 7.4, 4% [w/v] SDS, 40% [v/v] glycerol, 10% [v/v] β-mercaptoethanol, 0.002% [w/v] bromphenol blue) were added. The samples (0.5–1.0 mg/mL of protein) were boiled for 5 min and 30 mL were loaded on 10% SDS-PAGE. The proteins were separated by electrophoresis and transferred to nitrocellulose membranes. Anti-GFP mouse IgG (clones 7.1 and 13.1; Roche Applied Science) was used at a dilution of 1:2,000 and it was visualized by incubation with goat anti-mouse IgG secondary antibodies conjugated to horseradish peroxidase (1:2,000) followed by a chemiluminescence reaction (Clarity Western ECL Substrate; Bio-rad). For the SS-TOM lines analysis, proteins were extracted in 2x SDS buffer (4% SDS, 125mM Tris pH 6.8, 20% glicerol, 0.01% bromophenol blue, 50 mM dithiothreitol [DTT]), using 10 μl of buffer per mg of plant tissues of Wt Col-0, transgenic lines 35S:SS-TOM and 35S:SS-TOM-Long-EXT. Two transgenic lines were analyzed. 10 μl of supernatant of each protein extract were run into a 12% polyacrylamide gel during one hour at 200 V, and then transferred to a PVDF membrane. PVDF was blocked with 5% milk in TBST (Tris-HCl 10 mM, pH 7,4, NaCl 150 mM, Tween-20 al 0,05%) for 1 hour at 4ºC and then washed four times during 15 min in TBST. An anti-RFP (A00682, GenScript) was used as primary antibody overnight at 4ºC. Four washes of 15 min each in TBST at room temperature and then it was incubated two hours with a secondary antibody anti-rabbit (goat) conjugated with alkaline phosphatase (A3687, Sigma), in a 1:2,500 dilution with TBST. Four washes of 15 min each in TBST at room temperature. Finally, 10 ml of alkaline phosphatase (100mM Tris-HCl pH 9.5, 100 mM NaCl, 3 mM MgCl2) containing 80 μl NBT (Sigma) (35mg/ml in70% DMSO and 30 μl de BCIP (Sigma) (50 mg/ml in 100% de DMSO) were used.

### Transmission electron microscopy of root hair cell walls

Seeds were germinated on 0.2x MS, 1% sucrose, 0.8% agar. Seven days after germination, seedlings were transferred to new 0.2x MS, 1% sucrose, 0.8% agar plates with or without 100 µM SHAM. After 4 additional days, 1-mm root segments with root hairs were fix in 2% glutaraldehyde in 0.1M cacodylate buffer pH7.4. Samples were rinsed in cacodylate buffer and post-fixed in 2% OsO4. After dehydration in ethanol and acetone, samples were infiltrated in Epon resin (Ted Pella, Redding, CA). Polymerization was performed at 60°C. Sections were stained with 2% uranyl acetate in 70% methanol followed by Reynold’s lead citrate (2.6% lead nitrate and 3.5% sodium citrate [pH 12.0]) and observed in a Tecnai 12 electron microscope. Quantitative analysis of cell wall thickness was performed using FIJI.

### Modeling and molecular docking between PRXs and EXTs

Modeling and molecular docking: cDNA sequences of PRXs were retrieved from TAIR (PRX01: AT1G05240, PRX36: AT3G50990, PRX44: AT4G26010, PRX64: AT5G42180, PRX73: AT5G67400) and NCBI Nucleotide DB (PRX24Gv:Vitis vinifera peroxidase 24, GvEP1, LOC100254434). Homology modeling was performed for all PRXs using modeller 9.14 (Sali et al. 1993), using the crystal structures 1PA2, 3HDL, 1QO4 and 1HCH as templates, available at the protein data bank. 100 structures where generated for each protein and the best scoring one (according to DOPE score) was picked. The *receptor* for the docking runs was generated by the prepare_receptor4 script from autodock suite, adding hydrogens and constructing bonds. Peptides based on the sequence PYYSPSPKVYYPPPSSYVYPPPPS were used, replacing proline by hydroxyproline, and/or adding *O*-Hyp glycosylation with up to four arabinoses per hydroxyproline in the fully glycosylated peptide and a galactose on the serine, as it is usual in plant *O*-Hyp https://www.ncbi.nlm.nih.gov/pmc/articles/PMC5045529/. Ligand starting structure was generated as the most stable structure by molecular dynamics (Velasquez et al. 2015a). All ligand bonds were set to be able to rotate. Docking was performed in two steps, using Autodock vina (Trott et al. 2010). First, an exploratory search over the whole protein surface (exhaustiveness 4) was done, followed by a more exhaustive one (exhaustiveness 8), reducing the search space to a 75×75×75 box centered over the most frequent binding site found in the former run.

### EXT conformational coarse-grained model

The use of coarse-grained (CG) molecular dynamics (MD) allowed collection of long timescale trajectories. System reduction is significant when compared to all atom models, approximately reducing on order of magnitude in particle number. In addition, a longer integration time step can be used. Protein residues and coarse grained solvent parameters correspond to the SIRAH model (Darré et al. 2015), while ad hoc specific glycan parameters were developed. The CG force field parameters developed correspond to arabinofuranose and galactopyranose (**Figure S5**). Triple helix systems were simulated both, in the non-glycosylated and fully *O*-glycosylated states, where all the hydroxyprolines are bound to a tetrasaccharide of arabinofuranoses, and specific serine residues contain one galactopyranose molecule. They were immersed in WT4 GC solvent box that was constructed to be 2 nm apart from the extensin fiber, and periodic boundary conditions were employed. Coarse grained ions were also included to achieve electroneutrality and 0.15 M ionic strength. All simulations were performed using the GROMACS MD package at constant temperature and pressure, using the Berendsen thermostat (respectively) and Parrinello-Rahman barostat (Parrinello and Rahman 1981), and a 10 fs time step. The obtained trajectories were analysed using the Mdtraj python package (McGibbon et al, 2015) and visualized with Visual Molecular Dynamics (VMD) 1.9.1 (Humphrey et al. 1996). Volume measurements were performed using a Convex Hull algorithm implemented in NumPy (Oliphant 2006), and average diameter calculations were derived from this quantity using simple geometric arguments.

## Acknowledgements

We thank Andres Rossi from the microscopy facility of FIL for his assistance. We thank ABRC (Ohio State University) for providing T-DNA lines seed lines. J.M.E. is Principal Investigator of the National Research Council (CONICET) from Argentina. This work was supported by grants from ANPCyT (PICT2016-0132 and PICT2017-0066) and a grant from ICGEB CRP/ARG16-03 to J.M.E. In addition, this research is also funded by Instituto Milenio iBio–Iniciativa Científica Milenio, MINECON to J.M.E. and NSF MCB grant 1614965 to M.S.O.

## Author Contribution

E.M and C.B performed most of the experiments and analysed the data. P.R. and C.D. analysed the peroxidase activity and performed phylogenetic analysis. J.W.M-E and M.H. analysed the Tyr-crosslinking on EXTs. A.A.A. and A.D.N performed the docking experiments and analysed this data. M.B. and L.C. perform the EXT modelling and analysed this data. M.F. and P.B generated the EXT reporter lines and performed the immune-blots analysis of SS-TOM and SS-TOM-Long-EXT lines. J.M.P., D.R.R.G., Y.d.C.R.G., S.M., and F.B.H. analysed the data. J.P., J.P-V., and M.S.O. performed the transmission electron microscopy analysis. J.M.E. designed research, analysed the data, supervised the project, and wrote the paper. All authors commented on the results and the manuscript. This manuscript has not been published and is not under consideration for publication elsewhere. All the authors have read the manuscript and have approved this submission.

## REFERENCES

Baumberger N, Ringli C, Keller B. (2001). The chimeric leucine rich repeat/extensin cell wall protein LRX1 is required for root hair morphogenesis in *Arabidopsis thaliana*. Genes & Development 15, 1128–1139.

Baumberger N, Steiner M, Ryser U, Keller B, Ringli C. (2003). Synergistic interaction of the two paralogous Arabidopsis genes LRX1 and LRX2 in cell wall formation during root hair development. Plant J. 35, 71–81.

Berendsen, H. J. C., Postma, J.P.M., van Gunsteren, W.F., DiNola, A., and Haak, J.R. (1984) Molecular dynamics with coupling to an external bath. J. Chem. Phys. 81, 3684. doi: 10.1063/1.448118.

Berman HM, John Westbrook, Zukang Feng, Gary Gilliland, Talapady N Bhat, Helge Weissig, Ilya N Shindyalov, and Philip E Bourne. The protein data bank. Nucleic Acids Research, 28(1):235–242, 2000.

Bernards, M.A., Fleming, W.D., Llewellyn, D.B., Priefer, R., Yang, X., Sabatino, A., and Plourde, G.L. (1999). Biochemical characterization of the suberization-associated anionic peroxidase of potato. Plant Physiol. 121: 135–146.

Bhave G, Cummings CF, Vanacore RM, Kumagai-Cresse C, Ero-Tolliver IA, Rafi M, Kang J-S, Pedchenko V, Fessler LI, Fessler JH, Hudson BG (2012) Peroxidasin forms sulfilimine chemical bonds using hypohalous acids in tissue genesis. Nat. Chem. Biol. 8:784–790.

Brady, J.D., Sadler, I.H., and Fry, S.C. (1996). Di-isodityrosine, a novel tetrametric derivative of tyrosine in plant cell wall proteins: a new potential cross-link. Biochem. J 315: 323–327.

Brady, J.D., Sadler, I.H., and Fry, S.C. (1998). Pulcherosine, an oxidatively coupled trimer of tyrosine in plant cell walls: its role in cross-link formation. Phytochemistry. 47(3):349–353.

Brown K.L, C.F. Cummings, R.M. Vanacore and B.G. Hudson. (2017). Building collagen IV smart scaffolds on the outside of cells. Protein Sci. 26:2151–2161.

Brownleader, M.D., Ahmed, N., Trevan, M., Chaplin, M.F., and Dey, P.M. (1995). Purification and partial characterization of Tomato extensin peroxidase. Plant Physiol. 109: 1115–1123

Cannon, M.C., Terneus, K., Hall, Q., Tan, L., Wang, Y., Wegenhart, B.L., Chen, L., Lamport, D.T., Chen, Y., and Kieliszewski, M.J. (2008). Self-assembly of the plant cell wall requires an extensin scaffold. Proc. Natl. Acad. Sci. USA. 105: 2226–2231.

Chen, Y., Dong, W., Tan, L., Held, M. A., & Kieliszewski, M. J. (2015). Arabinosylation plays a crucial role in extensin cross-linking in vitro. Biochemistry insights, 8, BCI–S31353.

Cosio C. & Dunand C. (2009). Specific functions of individual class III peroxidases genes. J. Exp. Bot. 62:391–408.

Cosio C., Ranocha Ph., Francoz E., Burlat V., Zheng Y., Perry S.E., Ripoll J.-J., Yanofsky M., Dunand C. (2017) The class III peroxidase PRX017 is a direct target of the MADS-box transcription factor 1 AGL15 and participates in lignified tissue formation. New Phytol. 213: 250–263.

Darré, L., Machado, M.R., Brandner, A.F., González, H.C., Ferreira, S., and Pantano, S. (2015). SIRAH: A structurally unbiased coarse-grained force field for proteins with aqueous solvation and longrange electrostatics. J. Chem. Theory Comput. 11(2):723–739.

Davey, C. A. and R. E. Fenna (1996). 2.3 Å resolution X-ray crystal structure of the bisubstrate analogue inhibitor salicylhydroxamic acid bound to human myeloperoxidase: a model for a prereaction complex with hydrogen peroxide. Biochemistry 35(33): 10967–10973.

Dong, W., Kieliszewski, M., and Held, M.A. (2015). Identification of the pI 4.6 extensin peroxidase from *Lycopersicon esculentum* using proteomics and reverse-genomics. Phytochem. 112:151–159.

Dunand, C., Crevecoeur, M., and Penel, C. (2007). Distribution of superoxide and hydrogen peroxide in Arabidopsis root and their influence on root development: possible interaction with peroxidases. New Phytol. 174:332–341.

Dynowski M, Schaaf G, Loque D, Moran O, Ludewig U (2008) Plant plasma membrane water channels conduct the signalling molecule H_2_O_2_. Biochem J 414: 53–56.

Egelund, J., Obel, N., Ulvskov, P., Geshi, N., Pauly, M., Bacic, A., and Larsen Petersen, B. (2007). Molecular characterization of two Arabidopsis thaliana glycosyltransferase mutants, rra1 and rra2, which have a reduced residual arabinose content in a polymer tightly associated with the cellulosic wall residue. Plant. Mol. Biol. 64:439–451.

Fabrice T, Vogler H, Draeger C, Munglani G, Gupta S, Herger A, Knox P, Grossniklaus U, Ringli C. (2018). LRX Proteins play a crucial role in pollen grain and pollen tube cell wall development. Plant Physiology 176: 1981–1992.

Francoz et al. (2019) Pectin demethylesterification generates platforms that anchor peroxidases to remodel plant cell wall domains. Developmental Cell 48, 1–16.

Hall, Q., and Cannon, M.C. (2002). The cell wall hydroxyproline-rich glycoprotein RSH is essential for normal embryo development in Arabidopsis. Plant Cell 14: 1161–1172.

Held, M. A., Tan, L., Kamyab, A., Hare, M., Shpak, E., and Kieliszewski, M. J. (2004). Di-isodityrosine is the intermolecular cross-link of isodityrosine-rich extensin analogs cross-linked in vitro. Journal of Biological Chemistry, 279(53), 55474–55482.

Herrero, J., Fernández-Pérez, F., Yebra, T., Novo-Uzal, E., Pomar, F., Pedreño, M.Á., Cuello, J., Guéra, A., Esteban-Carrasco, A., and Zapata, J.M. (2013). Bioinformatic and functional characterization of the basic peroxidase 72 from Arabidopsis thaliana involved in lignin biosynthesis. Planta 237: 1599–1612.

Hooijmaijers C, Rhee JY, Kwak KJ, Chung GC, Horie T, Katsuhara M, Kang H (2012) Hydrogen peroxide permeability of plasma membrane aquaporins of *Arabidopsis thaliana*. J Plant Res 125: 147–153.

Humphrey, W., Dalke, A., and Schulten, K. (1996). VMD: Visual molecular dynamics. J. Molec. Graphics 14:33–38

Ikeda-Saito, M., D. A. Shelley, L. Lu, K. Booth, W. Caughey and S. Kimura (1991). Salicylhydroxamic acid inhibits myeloperoxidase activity. Journal of Biological Chemistry 266(6): 3611–3616

Ishiwata A, Kaeothip S, Takeda Y, Ito Y (2014) Synthesis of the highly glycosylated hydrophilic motif of extensins. Angew Chem Int Ed Engl 53: 9812–9816

Jackson, P.A., Galinha, C.I., Pereira, C.S., Fortunato, A., Soares, N.C., Amâncio, S.B., and Pinto Ricardo, C.P. (2001). Rapid deposition of extensin during the elicitation of grapevine callus cultures is specifically catalyzed by a 40-kilodalton peroxidase. Plant Physiol. 127: 1065–1076.

Jacobowitz, J.R, Doyle W.C., and Weng, J-K. (2019). PRX9 and PRX40 are extensin peroxidases essential for maintaining tapetum and microspore cell wall integrity during Arabidopsis anther development. Plant Cell 31: 848–86.

Kim D Pruitt, Tatiana Tatusova, and Donna R Maglott. Ncbi reference sequences (refseq): a curated non-redundant sequence database of genomes, transcripts and proteins. Nucleic Acids Research, 35(suppl 1):D61–D65, 2006.

Kumar, S., Stecher G., Tamura K. (2016) MEGA7: molecular evolutionaary genetic analysis version 7.0 for bigger datasets. Mol. Biol. Evol. 33(7): 1870–1874.

Kunieda, T., Shimada, T., Kondo, M., Nishimura, M., Nishitani, K., and Hara-Nishimura, I. (2013). Spatiotemporal secretion of PEROXIDASE36 is required for seed coat mucilage extrusion in Arabidopsis. Plant Cell 25: 1355–1367.

Lamport, D.T.A., Kieliszewski, M.J., Chen, Y., and Cannon, M.C. (2011). Role of the extensin superfamily in primary cell wall architecture. Plant Physiol. 156: 11–19.

Lee, Y., Rubio, M.C., Alassimone, J., and Geldner, N. (2013). A mechanism for localized lignin deposition in the endodermis. Cell 153: 402–412.

Mangano S. et al. (2017). The molecular link between auxin and ROS-controlled root hair growth. Proc. Natl. Acad. Sci. U.S.A. 114(20):5289–5294.

Marzol E, Borassi C, Bringas M, Sede A, Rodríguez Garcia DR, Capece L, Estevez JM. (2018). Filling the Gaps to Solve the Extensin Puzzle. Mol Plant. 11(5):645–658.

McGibbon R.T, Beauchamp K.A., Harrigan M.P., Klein C., Swails, J.M., Hernández C.X., Schwantes C.R., Wang, L.P., Lane T.J., Pande V.S. (2015) MDTraj: A Modern Open Library for the Analysis of Molecular Dynamics Trajectories. Biophysical Journal, 109(8): 1528–1532.

Møller, S.R., et al. (2017). Identification and evolution of a plant cell wall specific glycoprotein glycosyl transferase, ExAD. Sci. Rep. 7: 45341.

Monshausen GB, Bibikova TN, Messerli MA, Shi C, Gilroy S (2007) Oscillations in extracellular pH and reactive oxygen species modulate tip growth of Arabidopsis root hairs. Proc Natl Acad Sci USA 104:20996–21001.

Mutwil, M., Klie, S., Tohge, T., Giorgi, F.M., Wilkins, O., Campbell, M.M., Fernie, A.R., Usadel, B., Nikoloski, Z. and Persson, S. (2011). PlaNet: combined sequence and expression comparisons across plant networks derived from seven species. Plant Cell, 23, 895–910.

Oliphant, TE. USA: Trelgol Publishing, (2006). A guide to NumPy. 376 pp.

Orman-Ligeza B, et al. (2016) RBOH-mediated ROS production facilitates lateral root emergence in Arabidopsis. Development 143:3328–3339

Owens NW, Stetefeld J, Lattová E, Schweizer F (2010) Contiguous *O*-galactosylation of 4(R)-hydroxy-l-proline residues forms very stable polyproline II helices. J Am Chem Soc 132: 5036–5042

Parrinello, M. and Rahman, A. (1981). Polymorphic transitions in single crystals: A new molecular dynamics method. J. of Applied Physics 52:7182. doi.org/10.1063/1.328693

Passardi, F., Penel, C., and Dunand, C. (2004). Performing the paradoxical: how plant peroxidases modify the cell wall. Trends Plant Sci. 9:534–540.

Pereira S.P., J.M.L. Ribeiro, A.D. Vatulescu, K. Findlay, A.J. MacDougall and P.A.P. Jackson. (2011) Extensin network formation in *Vitis vinifera* callus cells is an essential and causal event in rapid and H_2_O_2_-induced reduction in primary cell wall hydration. BMC Plant Biology 111:106–121.

Price, N.J., Pinheiro, C., Soares, C.M., Ashford, D.A., Ricardo, C.P., and Jackson, P.A. (2003). A biochemical and molecular characterization of LEP1, an extensin peroxidase from lupin. J. Biol. Chem. 278:41389–41399.

Ringli C. (2010). The hydroxyproline-rich glycoprotein domain of the Arabidopsis LRX1 requires Tyr for function but not for insolubilization in the cell wall. Plant J. 63, 662–669.

Rodrigues, O. et al. (2017). Aquaporins facilitate hydrogen peroxide entry into guard cells to mediate ABA- and pathogen-triggered stomatal closure. Proc Natl Acad Sci USA 114, 9200–9205, doi:10.1073/pnas.1704754114.

Saito, F., Suyama, A., Oka, T., Yoko-o, T., Matsuoka, K., Jigami, Y., and Shimma, Y. (2014). Identification of novel peptidyl serine O-galactosyltransferase gene family in plants. J. Biol. Chem. 30:20405–20420.

Saitou N, Nei M. (1987). The neighbor-joining method: a new method for reconstruction of phylogenetic trees. Mol Biol Evol 4:406–25.

Šali A., and Tom L Blundell. (1993). Comparative protein modelling by satisfaction of spatial restraints. J. of Molecular biology, 234(3):779–815.

Savelli B., Li Q., Webber M., Jemmat A.M., Robitaille A., Zamocky M., Mathe C., Dunand C. (2019). RedoxiBase a database for ROS homeostasis regulated proteins. Redox Biol. doi.org/10.1016/j.redox.2019.101247.

Schnabelrauch, L.S., Kieliszewski, M., Upham, B.L., Alizedeh, H., and Lamport, D. (1996). Isolation of pl 4.6 extensin peroxidase from tomato cell suspension cultures and identification of Val-Tyr-Lys as putative intermolecular cross-link site. Plant J. 9:477–489.

Sede, A.R., Borassi, C., Wengier, D.L., Mecchia, M.A., Estevez, J.M., and Muschietti, J.P. (2018). Arabidopsis pollen extensins LRX are required for cell wall integrity during pollen tube growth. FEBS Lett. 592, 233–243.

Shen Y., Rosendale M., Campbell R.E., D. Perrais (2014). pHuji, apH-sensitive red fluorescent protein for imaging of exo- and endocytosis. J.Cell Biol. 207 (3):419–432.

Smith, A.T., Santama, N., Dacey, S., Edwards, M., Bray, R.C., Thorneley, R.N., and Burke, J.F. (1990). Expression of a synthetic gene for horseradish peroxidase C in *Escherichia coli* and folding and activation of the recombinant enzyme with Ca2+ and heme. J. Biol. Chem. 265: 13335–13343.

Stafstrom, J.P., and Staehelin, L.A. (1986). The role of carbohydrate in maintaining extensin in an extended conformation. Plant Physiol. 81:242–246.

Stoddard, A and Rolland, V. (2018). I see the light!. Fluorescent proteins suitable for the cell wall/apoplat targeeting in *Nicotiana benthamiana* leaves. Plant Direct. doi.org/10.1002/pld3.112

Strasser, R. (2016). Plant protein glycosylation. Glycobiology. 26(9):926–939.

Stratford, S., et al. (2001). A leucine-rich repeat region is conserved in pollen extensin-like (Pex) proteins in monocots and dicots. Plant Mol. Biol. 46:43–56.

Takeda S, et al. (2008) Local positive feedback regulation determines cell shape in root hair cells. Science 319:1241–1244.

Trott O., and Arthur J Olson. (2010). Autodock vina: improving the speed and accuracy of docking with a new scoring function, efficient optimization, and multithreading. Journal of Computational Chem. 31(2):455–461.

Vanacore R, Ham AJ, Voehler M, Sanders CR, Conrads TP, Veenstra TD, Sharpless KB, Dawson PE, Hudson BG (2009) A sulfilimine bond identified in collagen IV. Science 325:1230–1234.

Velasquez, S.M., Ricardi, M.M., Dorosz, J.G., Fernandez, P.V., Nadra, A.D., Pol-Fachin, L., Egelund, J., Gille, S., Ciancia, M., Verli, H., et al. (2011). O-glycosylated cell wall extensins are essential in root hair growth. Science 332:1401–1403.

Velasquez M, Salter JS, Dorosz JG, Petersen BL, Estevez JM. (2012). Recent advances on the posttranslational modifications of EXTs and their roles in plant cell walls. Frontiers in plant science 3, 93.

Velasquez SM, Marzol E, Borassi C, Pol-Fachin L, Ricardi MM, Mangano S, Denita JS, Salgado SJ, Gloazzo DJ, Marcus SE. (2015a). Low sugar is not always good: Impact of specific O-glycan defects on tip growth in Arabidopsis. Plant Physiology 168, 808–813.

Velasquez, S.M., Ricardi, M.M., Poulsen, C.P., Oikawa, A., Dilokpimol, A., Halim, A., Mangano, S., Denita-Juarez, S.P., Marzol, E., Salgoda Salter, J.D., et al. (2015b). Complex regulation of prolyl-4-hydroxylases impacts root hair expansion. Mol. Plant 8:734–746.

Wang, X., Wang, K., Yin, G., Liu, X., Liu, M., Cao, N., Duan, Y., Gao, H., Wang, W., Ge, W., et al. (2018). Pollen-expressed leucine-rich repeat extensins are essential for pollen germination and growth. Plant Physiol. 176, 1993–2006.

Wojtaszek, P., Trethowan, J., and Bolwell, G.P. (1997). Reconstitution in vitro of the components and conditions required for the oxidative cross-linking of extracellular proteins in French bean (Phaseolus vulgaris L.). FEBS Lett. 405:95–98.

Yi K, Menand B, Bell E, Dolan L. (2010). A basic helix-loop-helix transcription factor controls cell growth and size in root hairs. Nature genetics 43: 264–267.

