## Supplemental materials for "Class III peroxidases PRX01, PRX44, and PRX73 potentially target extensins during root hair growth in *Arabidopsis thaliana*"

The following Supporting Information is available for this article:
Supplementary Figures S1-S7
Supplementary Table S1-S2
Supplementary References 6

**Table S1**. Volume and average diameter measurements of EXTs from molecular dynamics simulations. These magnitudes were measured in the fully extended EXT system. Values shown are the mean ± standard deviation.

| | **Non-glycosylated EXT** | ***O*-glycosylated EXT** | ***O*-glycosylated EXT (protein only)** |
|---|---|---|---|
| Volume /$nm^3$ | 280 ± 30 | 1080 ± 50 | 450 ± 30 |
| EXT length /nm | 65 ± 2 | 70 ± 2 | 70 ± 2 |
| Average diameter /nm | 2.4 | 4.5 | 2.9 |
| Distance Tyr-Tyr | | | |
| 0-5 Å | 20% | 25% | - |
| 5-7 Å | 55% | 65% | - |

**Table S2**. Mutants and transgenic lines generated and used in this study.

| Line name | Gene construct/ mutant lines code | References |
|---|---|---|
| **SS-TOM-Long-EXT #2** | 35Spromoter::SS-TOMATO-longEXT | This study |
| **SS-TOM-Long-EXT #7** | 35Spromoter::SS-TOMATO-longEXT | This study |
| **SS-TOM #2** | 35Spromoter::SS-TOMATO | This study |
| **SS-TOM #5** | **3**5Spromoter::SS-TOMATO | This study |
| **PRX01::GFP** | PRX01promoter::GFP | This study |
| **PRX44::GFP** | PRX44promoter::GFP | This study |
| **PRX73::GFP** | PRX73promoter::GFP | This study |
| **35S::PRX01-1** | 35Spromoter::PRX01-GFP | This study |
| **35S::PRX01-2** | 35Spromoter::PRX01-GFP | This study |
| **35S::PRX44-3** | 35Spromoter::PRX44-GFP | This study |
| **35S::PRX44-4** | 35Spromoter::PRX44-GFP | This study |
| **35S::PRX73-2** | 35Spromoter::PRX73-GFP | This study |
| **35S::PR7X3-3** | 35Spromoter::PRX73-GFP | This study |
| *prx01,44,73* | *prx01-2* (SALK_103597)<br>*prx44-2* (SALK_057222)<br>*prx73-3* (SALK_009296) | This study |
| *sgt1-1 rra3* | *sgt1-1* (SALK_054682)<br>*rra3* (GABI_233B05) | Velasquez et al. 2015a |
| *p4h5 sergt1-1* | *p4h5* (SALK_152869)<br>*sgt1-1* (SALK_054682) | Velasquez et al. 2015a |

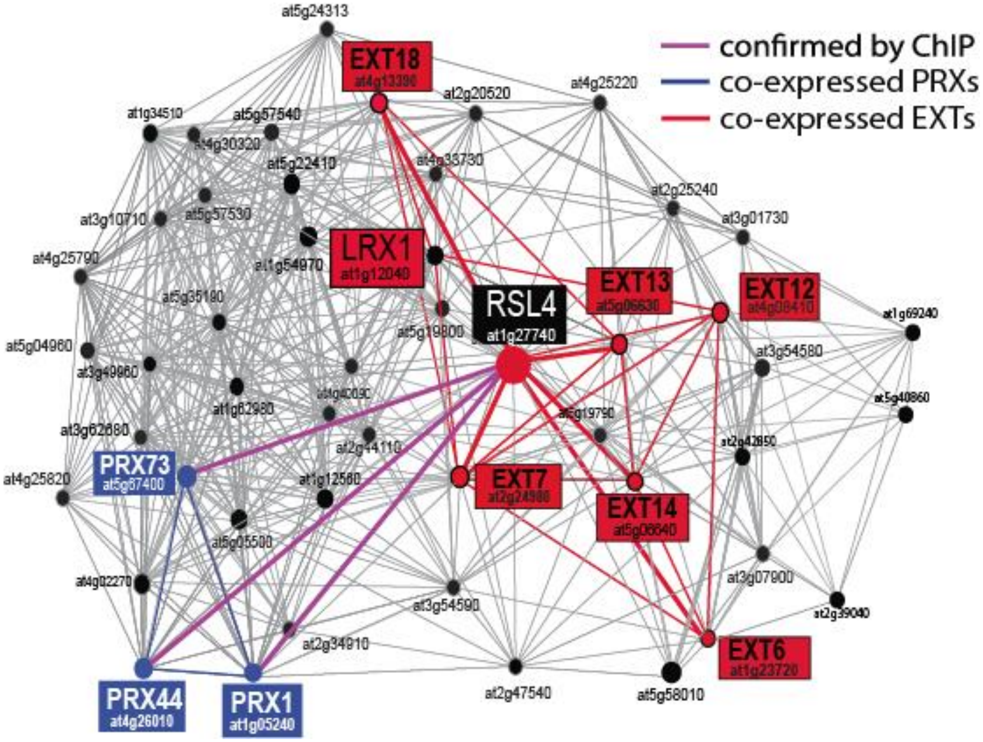

**Figure S1**. **Root hair transcriptional co-expression network.** PRXs, EXTs, and the transcriptional regulator RSL4 are highlighted. RSL4 was used as gene bait to narrow down the number of co-expressed genes. Transcriptional connections between RSL4 (in black), EXTs (in red) and PRXs genes (in blue). The co-expression network was identified from PLaNet (http://aranet.mpimp-golm.mpg.de/aranet) (Mutwil et al., 2011) and trimmed to facilitate readability. Chromatin immunoprecipitation assay (ChiP) showed that RSL4 binds and controls the expression of PRX01,44,73 (Mangano et al. 2017). Figure modified from Marzol et al. (2018).

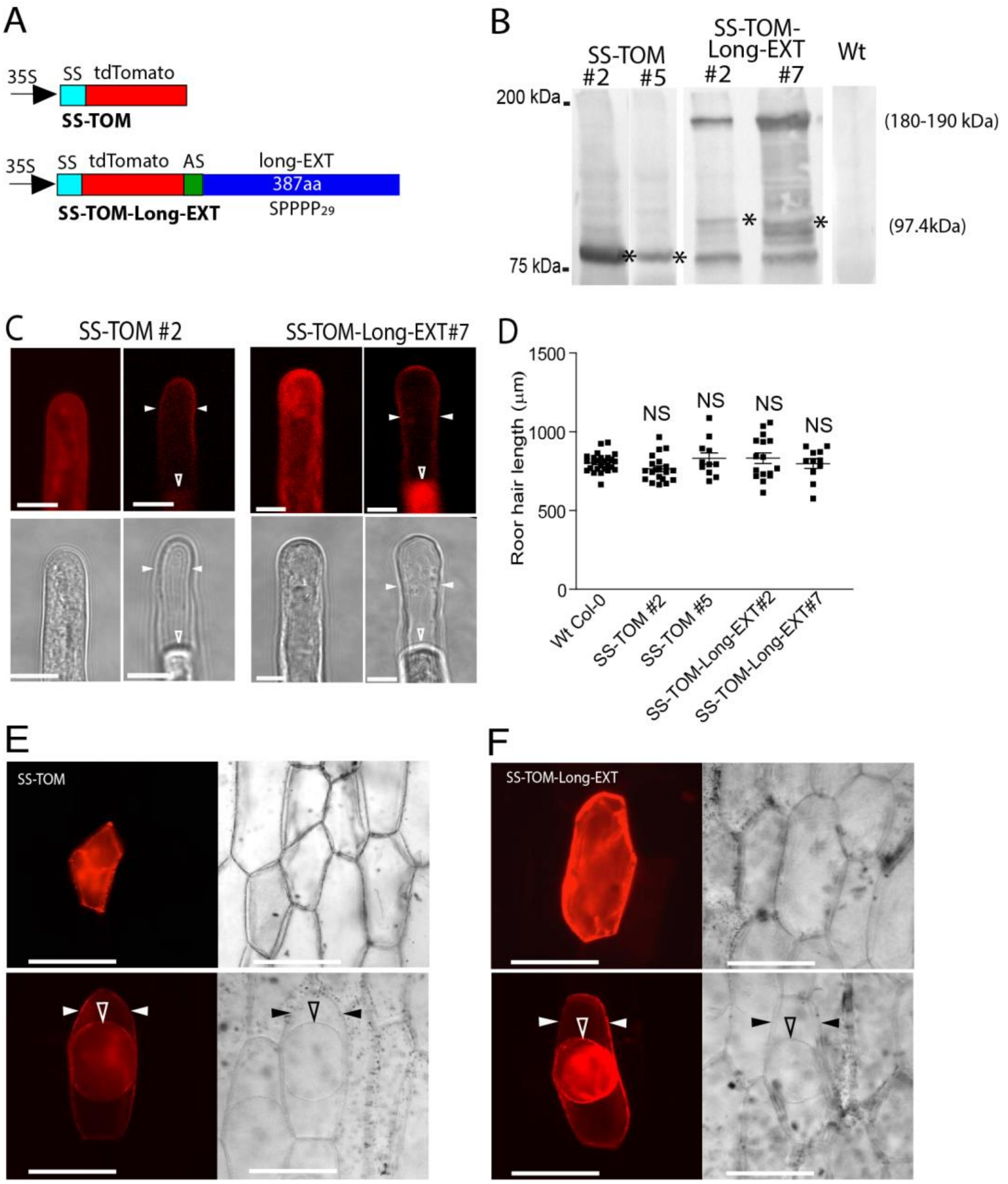

**Figure S2. EXT reporter is targeted to root hair and onion epidermal cell walls.**

(**A**) Schematic diagrams of SS-TOM and SS-TOM-Long-EXT constructs expressed under the control of the 35S promoter in Arabidopsis plants. SS, tomato polygalacturonase signal sequence. AS, Ala-spacer, 6 alanines between tdTomato (TOM) and Long-EXT domain. Long-EXT, C-terminal 405 amino acids of *SlPEX1*, which includes only two tyrosines at the C-terminus.

(**B**) Expression and detection of EXT-reporters in homozygous *Arabidopsis* lines. Lines expressing detectable levels of SS-TOM (lines #2 and #5) and SS-TOM-Long-EXT (lines #2 and #7) showed similarly unexpectedly large sizes of protein on the Western blot, with bands much larger than the predicted size of the unmodified TOM tagged Long-EXT protein. The predicted molecular size for SS-TOM protein is around 54.2 kDa and for SS-TOM-Long-EXT is around 97.4 kDa are indicated with asterisks (*). All the gel blots shown are part of the same run.

(**C**) Expression of SS-TOM and SS-TOM-Long-EXT reporters in *Arabidopsis* root hair cells. (Top) Fluorescence images showing cell-wall localization of the reporter protein: left, cells with normal cytoplasm; right, cells after plasmolysis in 1 M NaCl. (Bottom) Bright-field images of the same cells shown above. Cell wall location is denoted with solid arrowheads while plasma membrane is indicated with an unfilled arrowhead. Scale bar = 10 µm.

(**D**) Root hair phenotype of Wt, SS-TOM (#2 and #5) and SS-TOM-Long-EXT (#2 and #7) (in Wt background) as box-plot. Horizontal lines show the means. NS = not significantly different, determined by one-way ANOVA.

(**E**) Onion epidermis expressing SS-TOM. (On the right) Bright field images. (On the bottom) cells were plasmolyzed in 1 M NaCl. Cell wall location is denoted with solid arrowheads while plasma membrane is indicated with an unfilled arrowhead. Scale bar = 100 µm. SS = Signal Peptide. TOM = Tomato fluorescent protein.

(**F**) Onion epidermis expressing SS-TOM-Long-EXT. (On the right) Bright fields images. (On the bottom) Cells were plasmolyzed in 1 M NaCl. Cell wall location is denoted with solid arrowheads while plasma membrane is indicated with an unfilled arrowhead. Scale bar = 100 µm. SS = Signal Peptide. TOM = Tomato fluorescent protein. Long-EXT = C-terminal 290 amino acids of *Solanum lycopersicum* (Sl) *SlPEX1*.

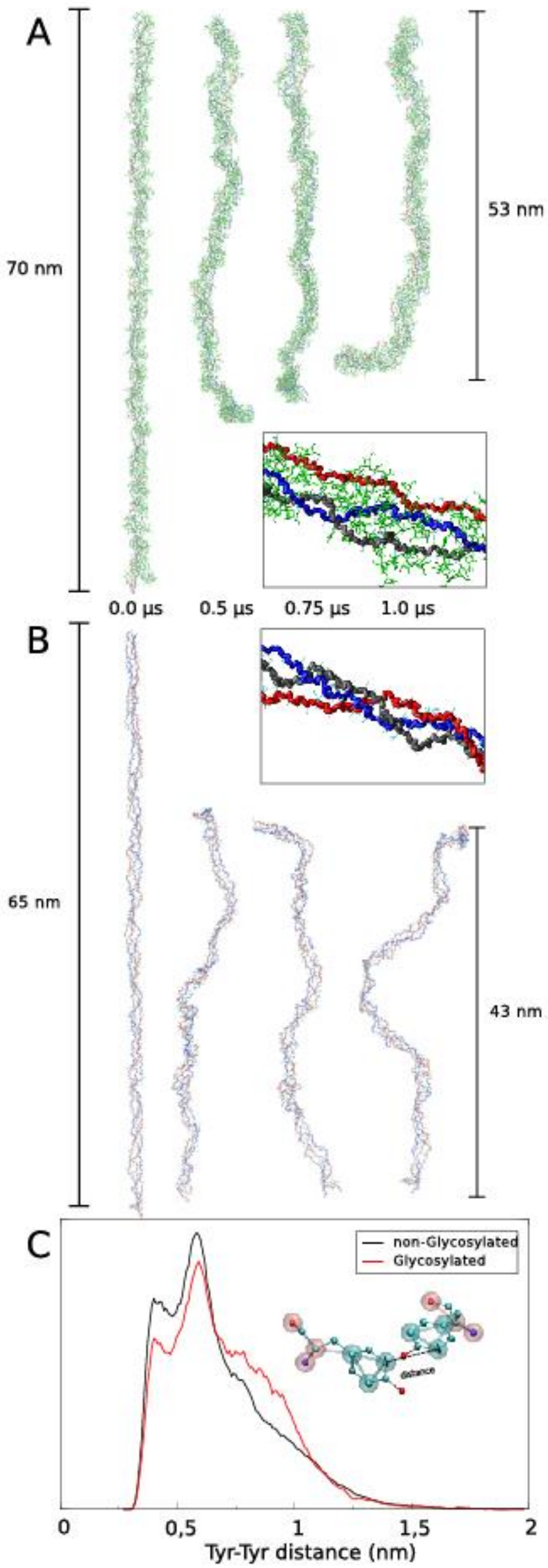
A
70 nm
53 nm
0.0 µs
0.5 µs
0.75 µs
1.0 µs
B
65 nm
43 nm
C
non-Glycosylated
Glycosylated
distance
0
0,5
1
1,5
2
Tyr-Tyr distance (nm)

69

**Figure S3. Effect of the *O*-glycosylation status on the triple helix EXT assembly and Tyr-Tyr distances.** Selected snapshots of the *O*-glycosylated (**A**) and non-glycosylated **(B)** triple EXT chain molecular dynamics (MD) trajectory, showing the stability of the triple helix during a 1 µs MD simulation. The results obtained in these simulations highlight the importance of the triple-helix EXT in overall protein stability, and especially the maintenance of the fibril-like structure. Peptide chains are depicted in red, black and blue and glycans in green. In these simulations the peptide chains are allowed to move freely inside the simulation box, without any restrictions. Insets: a snapshot of a 25 amino acid portion of the triple helix taken from the CG MD trajectory.

(**C**) Distribution of the Tyr-Tyr distances along the MD simulations for the glycosylated (red line) and non-glycosylated (black line) states. Distances are measured between the C-*ortho*s of each Tyr residue (as depicted in the inset of the figure).

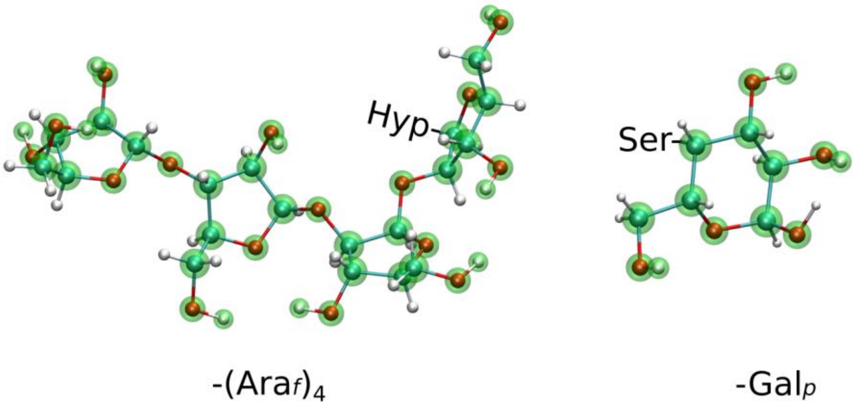

**Figure S4**. **Coarse-grained models for tetra-arabinofuranose and galactopyranose moieties (diffuse green spheres) superimposed on all atom description (solid spheres connected by tubes)**. Interaction points or beads are located in the position of the heavy atoms and hydrogens corresponding to hydroxyl groups. The latter were included due to the importance of directional hydrogen bond-like interactions in this class of molecules. Mass and Lennard-Jones parameters of CG beads and spring constants for intramolecular interactions were chosen to be compatible with the SIRAH coarse-grained model (parameters are available upon request).

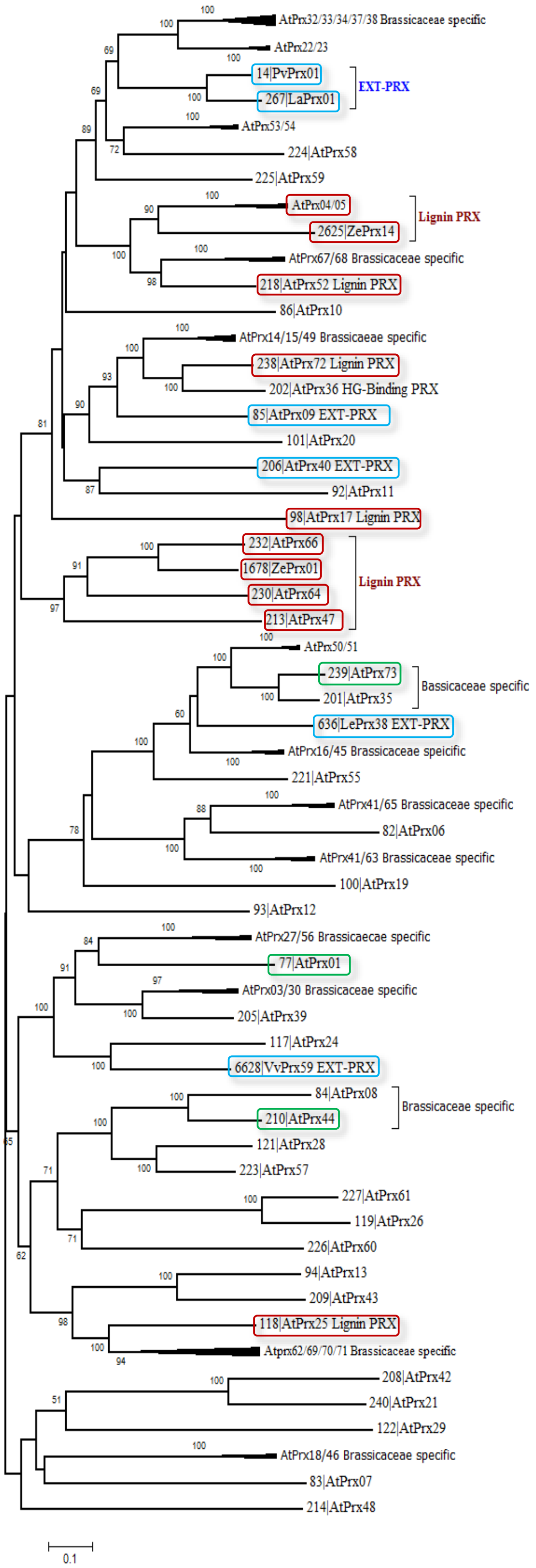
AtPrx32/33/34/37/38 Brassicaceae specific
AtPrx22/23
14|PvPrx01
267|LaPrx01
EXT-PRX
AtPrx53/54
224|AtPrx58
225|AtPrx59
AtPrx04/05
2625|ZePrx14
Lignin PRX
AtPrx67/68 Brassicaceae specific
218|AtPrx52 Lignin PRX
86|AtPrx10
AtPrx14/15/49 Brassicaceae specific
238|AtPrx72 Lignin PRX
202|AtPrx36 HG-Binding PRX
85|AtPrx09 EXT-PRX
101|AtPrx20
206|AtPrx40 EXT-PRX
92|AtPrx11
98|AtPrx17 Lignin PRX
232|AtPrx66
1678|ZePrx01
230|AtPrx64
213|AtPrx47
Lignin PRX
AtPrx50/51
239|AtPrx73
201|AtPrx35
Bassicaceae specific
636|LePrx38 EXT-PRX
AtPrx16/45 Brassicaceae specific
221|AtPrx55
AtPrx41/65 Brassicaceae specific
82|AtPrx06
AtPrx41/63 Brassicaceae specific
100|AtPrx19
93|AtPrx12
AtPrx27/56 Brassicaecae specific
77|AtPrx01
AtPrx03/30 Brassicaceae specific
205|AtPrx39
117|AtPrx24
6628|VvPrx59 EXT-PRX
84|AtPrx08
210|AtPrx44
Brassicaceae specific
121|AtPrx28
223|AtPrx57
227|AtPrx61
119|AtPrx26
226|AtPrx60
94|AtPrx13
209|AtPrx43
118|AtPrx25 Lignin PRX
Atprx62/69/70/71 Brassicaceae specific
208|AtPrx42
240|AtPrx21
122|AtPrx29
AtPrx18/46 Brassicaceae specific
83|AtPrx07
214|AtPrx48
0.1

**Figure S5. Evolutionary relationships of CIII Prx with putative extensin cross-linking activity.** The 73 Class-III Peroxidase protein sequences from *A. thaliana*, two putative Lignin PRX from *Zinnia elegans* and four putative EXT-PRX from *Lupinus album*, *Solanum lycopersicum*, *Phaseolus vulgaris* and *Vitis vinifera*, have been aligned with ClustalW and the tree constructed using the Neighbor-Joining method (Saitou and Nei, 1987). The analyses were conducted in MEGA7 (Kumar, 2016). Class-III PRXs with putative EXT-PRX activity or Lignin PRX activity were surrounded in blue and red respectively. The three Class-III PRXs studied in the present work were surrounded in green. Branches of the tree with sequences resulting from duplication events only detected in Brassicaceae have collapsed to simplify the tree topology.

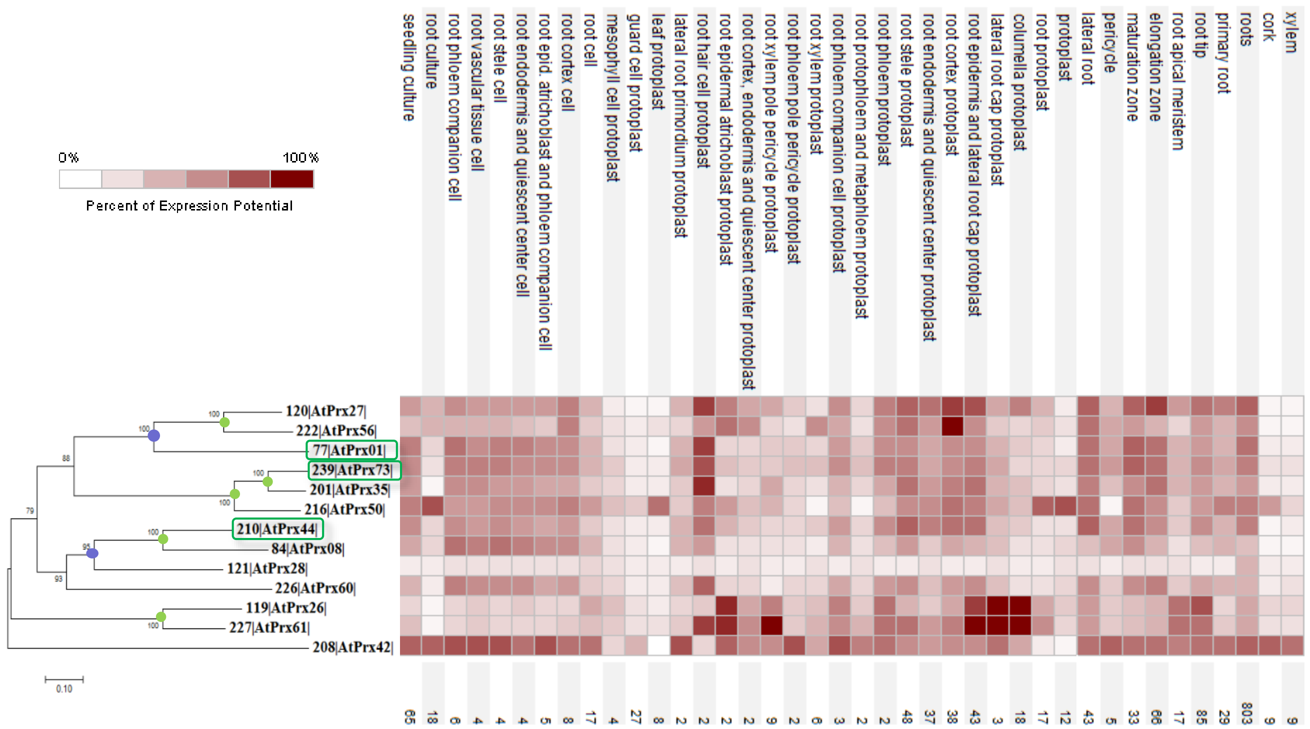

**Figure S6. Evolutionary and root tissue expression relationships of putative EXT-PRXs in Arabidopsis.**

The evolutionary history of Class III PRXs in root hair associated has been conducted with 15 CIII PRX protein sequences of *A. thaliana*. They have been aligned with ClustalW and the tree constructed using the Neighbor-Joining method (Saitou and Nei, 1987). The analyses were conducted in MEGA7 (Kumar, 2016). The sequences of AtPRX01/02 and AtPRX50/51 are either identical or very close, leading to only two independent expression profiles from Affymetrix data. A brown (max value)-to-cream (min value) heatmap was drawn from 105 anatomical parts from data selection available from AT_AFFY_ATH1-0 and plotted in front of each branch and using Genevestigator. The 3 CIII PRX studied in the present work were surrounded in green. Green and blue circles correspond to duplication posterior to Brassicaceae and to Dicotyledons emergences respectively. All protein sequences are available using their ID number (http://peroxibase.toulouse.inra.fr, (Savelli et al., 2019). Numbers indicate the amount of dataset available for each tissue.

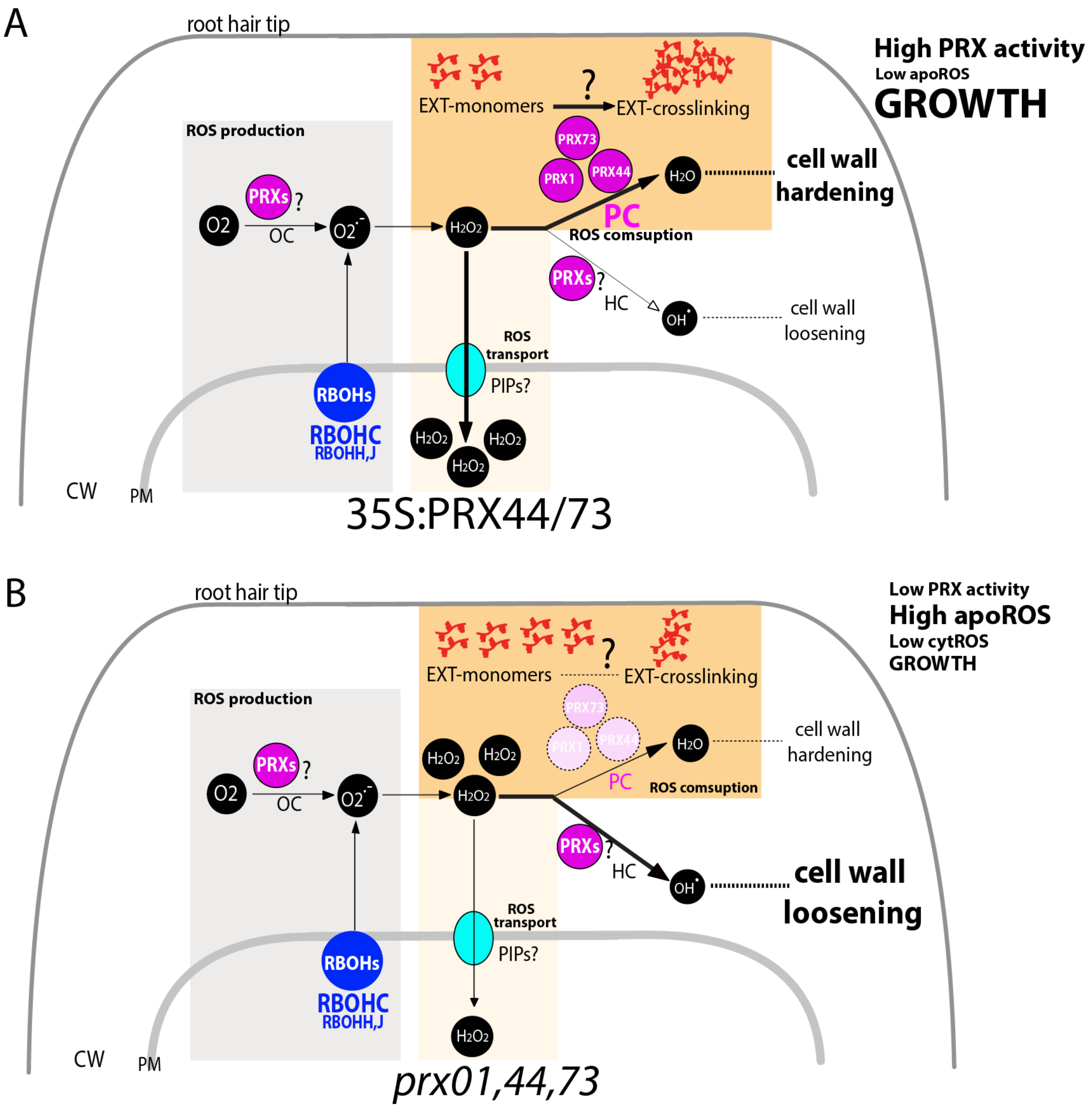

**Figure S7. Current model of PRX01, PRX44, and PRX73 function linking ROS homeostasis and EXT crosslinking in the root hair cell walls.** ROS homeostasis in the apoplast includes ROS production, ROS consumption, and ROS transport to the cytoplasm. ROS as superoxide ion ($O2^-$) is primarily produced by RBOHC and by RBOHH,J [based on Mangano et al. (2017)] and, possibly, by unknown PRXs in the root hair tip, and then dismutated to hydrogen peroxide ($H_2O_2$). (**A**) It is proposed that high levels of PRX44 or PRX73 during the peroxidative cycle (PC) (as found in 35S:PRX44-GFP and 35S:PRX73-GFP lines) might trigger a high consumption of apoplastic $H_2O_2$ (low $_{apo}$ROS) with a concomitant cell wall hardening. (**B**) Under low levels of all three PRXs (as found in the triple mutant *prx01,44,73*), most of the $H_2O_2$ produced accumulates in the apoplast (high $_{apo}$ROS) triggering a cessation of cell expansion. A portion of apoplastic $H_2O_2$ might be transported into the cytoplasm by aquaporins (PIPs). The balance between cell wall hardening and cell wall loosening processes might be compromised, affecting polar root hair growth. CW = cell wall; PM = plasma membrane; HC = hydroxylic cycle; OC = oxidative cycle. PIP= Plasma membrane Intrinsic Proteins (aquaporins).

### SUPPLEMENTARY REFERENCES

Kumar, S., Stecher G., Tamura K. (2016) MEGA7: molecular evolutionaary genetic analysis version 7.0 for bigger datasets. *Mol. Biol. Evol.* 33(7): 1870-1874.

Mangano S. *et al.* (2017). The molecular link between auxin and ROS-controlled root hair growth. *Proc. Natl. Acad. Sci. U.S.A*. 114(20):5289-5294.

Marzol E, Borassi C, Bringas M, Sede A, Rodríguez Garcia DR, Capece L, Estevez JM. (2018). Filling the Gaps to Solve the Extensin Puzzle. *Mol Plant*. 11(5):645-658.

Mutwil, M., Klie, S., Tohge, T., Giorgi, F.M., Wilkins, O., Campbell, M.M., Fernie, A.R., Usadel, B., Nikoloski, Z. and Persson, S. (2011). PlaNet: combined sequence and expression comparisons across plant networks derived from seven species. *Plant Cell*, 23, 895– 910.

Saitou N, Nei M. (1987). The neighbor-joining method: a newmethod for reconstruction of phylogenetic trees. Mol Biol Evol4:406–25.

Savelli B., Li Q., Webber M., Jemmat A.M., Robitaille A., Zamocky M., Mathe C., Dunand C. (2019). RedoxiBase a database for ROS homeostasis regulated proteins. *Redox Biol*. doi.org/10.1016/j.redox.2019.101247.

Velasquez SM, Marzol E, Borassi C, Pol-Fachin L, Ricardi MM, Mangano S, Denita JS, Salgado SJ, Gloazzo DJ, Marcus SE. (2015a). Low sugar is not always good: Impact of specific O-glycan defects on tip growth in Arabidopsis. *Plant Physiology* 168, 808–813.
